# Feedback Valence, Prediction Error, and the Limits of Computational Attribution in Symptom-Related EEG

**DOI:** 10.64898/2026.11.04.762513

**Authors:** Vaibhav Sinha, Vikram Ravindra

**Author notes:** **Correspondence**: Vaibhav Sinha.

## Abstract

Feedback-locked electroencephalography (EEG) responses are often interpreted using model- derived prediction errors (PEs), although signed PE can be strongly coupled to feedback valence in binary-outcome tasks. We revisited a published anxiety-related EEG association using two open probabilistic-learning datasets to examine measurement reliability, predictor separability, and participant-level recovery. In the primary dataset, greater trait anxiety was associated with a larger negative-minus-positive FCz theta response among 46 participants with elevated depressive symptoms after adjustment for depression severity. The association also differed between the sampled depression-symptom groups. The feedback contrast showed useful within-session reliability, but matched delta and P3-minus-N2 analyses did not establish theta specificity. Within participants, feedback valence and signed PE were highly correlated (*r* = *−*0.964). Exact- design simulations showed that population-level model discrimination could succeed despite poor recovery of individual neural slopes. Participant-specific learning-model refits recovered overall signed PE more consistently than the within-valence variation needed to distinguish it from feedback valence. An independent dataset reproduced the FCz feedback contrast in untreated and placebo cohorts but lacked trait-anxiety scores. Adding signed PE to valence yielded a small held-out predictive improvement under one reinforcement-learning specification, with limited support across sessions and no comparable support under the dual-rate model or in placebo. Together, these findings support a symptom-related feedback EEG association while constraining its theta-specific and participant-level computational interpretation.

## 1 Introduction

Feedback-related electroencephalography (EEG) signals are often interpreted in terms of reinforcement- learning computations, particularly prediction error (PE). Outcome evaluation is expressed in several frontocentral responses. Miltner et al. (1997) characterized the feedback-related negativity (FRN), and Proudfit (2015) reviewed the reward positivity (RewP) as a reward-sensitive event-related response. Theta- and delta-band activity provide complementary time–frequency measures of feed- back processing (Bernat et al., 2015; Cavanagh et al., 2010; Williams et al., 2021). Reinforcement learning accounts connect deviations between expected and obtained outcomes to trialwise neural signals (Cavanagh et al., 2010; Holroyd and Coles, 2002; Sambrook and Goslin, 2015). Frontal theta also participates in cognitive-control and performance-monitoring processes (Cavanagh and Frank, 2014). However, a reliable feedback-locked response does not by itself identify the computation represented by that response.

This problem is relevant to computational studies of depression and anxiety. Punishment sensitivity is associated with both symptom domains, whereas reduced reward sensitivity appears more selectively related to depression (Katz et al., 2020). Reinforcement learning findings in mood and anxiety disorders vary across tasks, parameters, and clinical populations (Pike and Robinson, 2022). Jiang et al. (2018) also associated trait anxiety with altered probabilistic feedback learning and feedback-locked electrophysiology. In the dataset motivating the present study, Cavanagh et al. (2019) reported that anxiety among participants with elevated depressive symptoms was associated with avoidance learning and stronger coupling between punishment-related EEG activity and negative prediction errors. The original result establishes a symptom association with an EEG signal expressed in fitted-PE terms, but leaves open whether that association depends on expectation-related variation beyond observed feedback valence and whether such variation can be estimated for individual participants. Other studies have reported nonuniform associations among depression, anxiety, and reward- or punishment-related EEG features (Grabowska et al., 2024; Rohlinger et al., 2026).

The interpretation of signed PE is constrained by the structure of binary-feedback tasks. On trial *t*, let *a_t_* denote the chosen action, *r_t_*the observed binary outcome, and *Q_t_*(*a_t_*) the action value assigned to that choice before feedback. Signed PE, denoted *δ_t_*, is

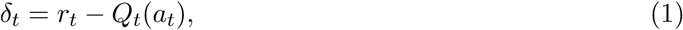

with *r_t_ ∈ {*0, 1*}* and *Q_t_*(*a_t_*) *∈* [0, 1]. On the 0–1 reward scale used here, positive feedback gives *δ_t_* = 1 *− Q_t_*(*a_t_*) and negative feedback gives *δ_t_* = *−Q_t_*(*a_t_*). Feedback valence determines the direction of signed PE, while the current action value determines its magnitude. Strong empirical correlation between valence and signed PE can arise even when the action values vary meaningfully across trials. When the two predictors carry little independent variation, a fitted neural association cannot be assigned cleanly to one of them from a single-predictor model. Nassar et al. (2019) showed that statistical context can alter the relation between feedback EEG and learning variables. Valence and expectancy-related information also need not occupy the same temporal or spectral components across paradigms (Stolz et al., 2023; Watts et al., 2017). Tasks designed to vary outcome dimensions more independently can separate overlapping PE processes (Hoy et al., 2021), but that separation cannot be assumed in a binary probabilistic-selection task (Zhang et al., 2020).

Individual-difference claims introduce a second requirement. A neural effect can be robust at the group level while providing limited dependable between-person variation. Clayson and Miller (2017) showed why reliability must be evaluated for the specific score, task, and analysis procedure. Difference scores can be less reliable than their constituent responses (Meyer et al., 2017), although reward- and error-related measures can achieve useful stability under suitable measurement conditions (Amir et al., 2023; Szenczy et al., 2025). Computational quantities face a related problem. Population- level model discrimination can succeed even when participant-specific parameters or neural slopes are recovered imprecisely (Wilson and Collins, 2019). Reinforcement learning parameters show variable test–retest reliability across tasks and parameterizations (Mkrtchian et al., 2023; Schaaf et al., 2024), and hierarchical estimation does not remove uncertainty that arises from weak information in the design (Katahira et al., 2024).

Independent data can test whether the neural measurement generalizes beyond the original sample. Such reproduction is narrower than replication of a symptom association or validation of a latent mechanism. High internal consistency can coexist with weak cross-task or symptom validity (Clayson et al., 2025), and reward-related EEG responses generalize more strongly across tasks that share similar structure (Lawhead et al., 2025). Clinical interpretation requires additional evidence beyond reproducibility of the neural measure (Clayson, 2025; Nielson et al., 2021; Simmatis et al., 2023).

Consequently, we revisited the symptom–EEG association reported by Cavanagh et al. (2019) to assess whether its interpretation as a participant-level PE effect is supported by the available data. In the primary dataset, we compared observed feedback valence with within-valence variation in fitted expected value and modeled STAI and BDI jointly within the elevated-BDI group. We tested whether the STAI–valence association differed between the two sampled BDI groups and assessed the feedback contrast’s reliability and temporal, spectral, and ERP characteristics. Exact-design simulations quantified cross-attribution between valence and PE, while refits to participant-specific simulated choices assessed recovery of the fitted learning variables. Additional simulations evaluated individual neural-slope recovery. An independent dataset then allowed us to reproduce the prespecified FCz response and test whether PE added predictive information beyond valence under alternative reinforcement-learning models.

## 2 Methods

### 2.1 Datasets, participants, and task

The primary analyses used OpenNeuro dataset ds003474, a public EEG dataset acquired during a probabilistic selection task (Cavanagh, 2021). Cavanagh et al. (2019) previously analyzed the release in relation to dimensional depression and anxiety symptoms. The dataset contains 122 participants, with Beck Depression Inventory (BDI) and trait scores from the State–Trait Anxiety Inventory (STAI) available for 121. The source study sampled two separated BDI groups. Its publication described the low-BDI criterion as BDI *<* 7, but the released 75-participant group includes one participant with BDI = 7. We retained the released group composition, defining the low-BDI group as BDI *≤* 7 (*N* = 75) and the elevated-BDI group as BDI *≥* 13 (*N* = 46). These labels refer to BDI score ranges and do not denote diagnostic groups.

During training, participants learned three probabilistic stimulus pairs, A/B (80/20), C/D (70/30), and E/F (60/40). Binary feedback was coded as reward (*r_t_* = 1) or nonreward (*r_t_* = 0). A later test phase presented novel pairings without feedback. Go performance was the proportion of eligible trials on which A was selected against stimuli other than B. NoGo performance was the proportion on which B was avoided when paired with stimuli other than A. The learning-bias score was NoGo minus Go, with positive values indicating relatively greater avoidance learning.

Independent analyses used OpenNeuro dataset ds004532 (Cavanagh and Frank, 2023). The release contains 110 participants and 137 recordings from a probabilistic selection paradigm that included a cabergoline manipulation (Cavanagh et al., 2014). STAI was not available. BDI and Behavioral Inhibition System (BIS) scores were available for a related-trait analysis. The measurement analyses used 77 untreated Study 1 recordings and 25 Study 2 placebo recordings with valid event structure before EEG trial-count eligibility. Cabergoline sessions and five early placebo-pilot recordings were excluded. Requiring at least 30 retained trials of each feedback valence yielded 70 Study 1 and 23 placebo sessions. Reliability analyses required at least 40 trials per valence and included 64 Study 1 and 20 placebo sessions. Additional task and event-reconstruction details are provided in Supplementary Methods.

### 2.2 EEG processing and neural measures

Primary-dataset EEG was sampled at 500 Hz. We read the continuous EEGLAB recordings distributed with ds003474 and reconstructed feedback events from the public task records. EEG preprocessing and feature extraction were implemented independently in MNE-Python (Gramfort et al., 2013), rather than using the source study’s cleaned epochs. Continuous data were notch filtered at 60 Hz, then zero-phase bandpass filtered from 0.5 to 100 Hz, and epoched from *−*2 to +2 s around feedback. Participant-relative channel and epoch metrics identified noisy or effectively flat data. Flagged scalp channels were interpolated before rereferencing to the average of mastoids M1 and M2. Independent component analysis (ICA) with FastICA (Hyvärinen and Oja, 2000) was used to remove components identified from horizontal and vertical electrooculogram (EOG) channels. The resulting preprocessed dataset contained 27,303 feedback epochs from all 122 participants. This count follows the exclusion of eight low-amplitude epochs by a post-ICA signal-integrity check. Screening criteria and implementation details are reported in Supplementary Methods, consistent with EEG reporting recommendations (Keil et al., 2014).

The recordings in the independent dataset differed in online reference, auxiliary-channel structure, and the scaling and conditioning of signals used for artifact detection. They used CPz as the online reference, which was not retained as a recorded channel. The single EOG channel in each recording had inconsistent absolute scaling across recordings, so no absolute EOG amplitude cutoff was used. The public continuous EEGLAB recordings contained 63 EEG channels and one EOG channel. We assigned standard 10–05 channel positions and applied average EEG rereferencing after spherical-spline interpolation. EEG and EOG channels were notch filtered at 60 Hz and zero-phase filtered from 0.5 to 40 Hz. Recording-relative robust statistics identified bad channels, with no more than 10 interpolated EEG channels permitted. FCz was not interpolated. Recordings in which it failed channel-quality screening were excluded from the analysis. Epochs were rejected using global EEG, FCz, and recording-relative EOG amplitude criteria. ICA was not used because decomposition conditioning varied across recordings and did not provide a common artifact-removal rule for this release. Artifact handling was tailored to the acquisition characteristics of each dataset. The definition of the neural measure was kept constant across datasets. Detailed criteria and the rationale for the dataset-specific artifact procedures are provided in Supplementary Methods.

The neural measure in both datasets was the FCz theta power from 4–8 Hz and 200–400 ms after feedback. The channel, frequency band, and latency followed the published analysis of ds003474 (Cavanagh et al., 2019). Fixing this region of interest before the present symptom analyses avoided selecting the neural target from the effects being tested. Time–frequency power was estimated with complex Morlet wavelets in MNE-Python (Gramfort et al., 2013). The wavelet parameters and corresponding time–frequency smoothing are specified in Supplementary Methods, following the reporting recommendations of Keil et al. (2022). Power was normalized in decibels (dB) to a participant- and frequency-specific baseline pooled across retained trials from *−*300 to *−*200 ms,

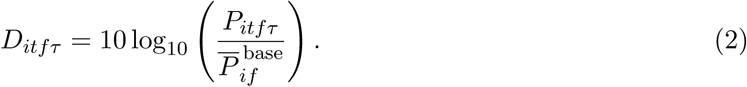

Here *D_itfτ_* is baseline-normalized power for participant *i*, trial *t*, frequency *f* , and time point *τ* . *P_itfτ_* is single-trial power at the same indices, and *P* ^base^ is the participant- and frequency-specific mean power in the pooled baseline interval. One theta value per trial was obtained by averaging *D_itfτ_* across 4–8 Hz and 200–400 ms. Trialwise theta was standardized within participant in the primary dataset and within participant-session in the independent dataset. The elevated-BDI symptom sample contributed 11,025 aligned trials, and the independent measurement sample contributed 19,312 trials from 93 sessions.

Additional neural measures were defined for reliability and electrophysiological characterization. Delta used 1–4-Hz power in the same latency interval. P3-minus-N2 was mean FCz voltage from 350–400 ms minus mean voltage from 250–300 ms after a 20-Hz low-pass filter and a *−*200 to 0 ms baseline. RewP used mean FCz voltage from 250–350 ms with the same filter and baseline.

### 2.3 Reinforcement learning models

Trialwise learning was modeled with Q-learning. Signed PE was defined by Equation 1, where *Q_t_*(*a_t_*) is the pre-feedback value of the chosen action, *r_t_* is the observed outcome, and *δ_t_*is the resulting signed PE. The chosen action value was updated as

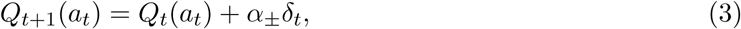

where *Q_t_*_+1_(*a_t_*) is the post-update value and *α_±_* is the learning rate selected according to the sign of *δ_t_*. Initial action values were *Q*_0_ = 0.5. A one-rate model used a common learning rate *α*. A separate-rate (dual-rate) model used *α*_+_ for nonnegative PEs and *α_−_* for negative PEs. The probability assigned to the observed choice among the two offered actions followed a binary softmax:

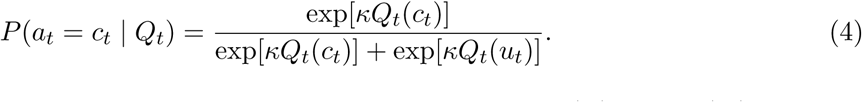

Here *c_t_*and *u_t_*denote the chosen and unchosen actions on trial *t*, and *Q_t_*(*c_t_*) and *Q_t_*(*u_t_*) are their estimated values before the feedback update. The inverse-temperature parameter *κ >* 0 controls choice sensitivity to the value difference. The choice probabilities approach one half as *κ* approaches zero and favor the higher-valued action increasingly as *κ* grows. The likelihood was evaluated using these pre-update values. Parameters were estimated separately for each participant by multistart maximum likelihood.

The primary-dataset decomposition used latent variables from the separate-rate model because that specification matched the prior EEG analysis (Cavanagh et al., 2019). The one-rate model served as a sensitivity analysis. In the independent dataset, the one-rate model was used as the primary computational specification, and the dual-rate model tested whether conclusions depended on the learning model. Optimizer settings and parameter bounds are given in Supplementary Methods.

Signed PE and feedback valence are structurally coupled in a binary-outcome task. Positive feedback gives *δ_t_* = 1 *− Q_t_*(*a_t_*), and negative feedback gives *δ_t_* = *−Q_t_*(*a_t_*). We used the conventional signed-PE regressor as a benchmark for the computational account without assuming that it identified an independent neural quantity. In the primary decomposition, positive- and negative-feedback PE magnitudes were centered within participant and feedback valence and entered alongside a direct negative-feedback indicator. A complementary positive-feedback parameterization used participant-centered chosen value. Since *|δ*^+^*|* = 1 *− Q_t_*(*a_t_*), positive-feedback PE magnitude and chosen value contain the same trialwise information with opposite sign.

### 2.4 Symptom and behavioral analyses

Feedback valence was coded *V_it_*= 1 for negative feedback and 0 for positive feedback, where *i* indexes participants and *t* indexes trials. Within the elevated-BDI sample, STAI and BDI were standardized across the 46 participants and entered together because the two symptom measures were correlated. We fitted a linear regression with participant fixed effects and interactions between feedback valence and the participant-level symptom measures:

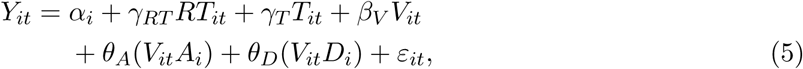

where *Y_it_* is within-participant standardized FCz theta, *A_i_* is standardized STAI, *D_i_* is standardized BDI, *RT_it_* is within-participant standardized response time, and *T_it_* is within-participant standardized trial progression. The participant-specific intercept *α_i_* implements the fixed effect for participant. The coefficients *γ_RT_* and *γ_T_* estimate the nuisance effects of response time and trial progression, *β_V_* estimates the feedback-valence effect at the reference symptom values, *θ_A_* and *θ_D_* estimate the STAI-by-valence and BDI-by-valence interactions, and *ε_it_* is the trial-level residual. Participant fixed effects absorb the symptom main effects. Response time and trial progression were included to account for within-participant variation associated with response latency and position in training; sensitivity analyses removed both covariates. The coefficient *θ_A_* tests whether STAI is associated with the negative-minus-positive feedback response after adjustment for BDI.

The decomposition model added valence-specific PE-magnitude regressors and their interactions with STAI and BDI. Chosen-value models assessed the equivalent positive-feedback parameterization, and one-rate latent variables assessed dependence on the learning model. Sensitivity analyses used raw-dB theta, an alternative within-trial baseline, removal of response-time and trial-progression covariates, and STAI residualized on BDI within the elevated-BDI group.

A separate group model compared the 75 low-BDI participants with the 46 elevated-BDI participants. STAI was expressed on its raw scale in 10-point units. Participant fixed effects were combined with valence-by-STAI, valence-by-group, and valence-by-STAI-by-group terms. The three-way coefficient tested whether STAI moderation of the feedback response differed between the sampled BDI groups.

For participant-level analyses, an adjusted negative-minus-positive FCz theta-gain score (hereafter, theta gain) was estimated separately for each elevated-BDI participant from a model containing feedback valence, response time, and trial progression. Partial Spearman correlations related theta gain to STAI while controlling BDI. The behavioral analysis related theta gain to NoGo-minus-Go performance. Participant-level inference used permutation tests and bootstrap confidence intervals. Detailed secondary symptom and behavioral analyses are reported in the supplement.

### 2.5 Reliability and electrophysiological characterization

Within-session consistency was estimated for theta gain, conventional signed-PE coupling, P3- minus-N2, delta, and RewP. Since the stability of difference measures can depend strongly on available observations (Clayson and Miller, 2017; Meyer et al., 2017), reliability was evaluated across several trial counts in addition to the full-session estimate. Eligible trials were divided by odd versus even order and by first versus second half. The main reliability statistic was the Spearman–Brown-corrected Pearson correlation, denoted *r*_SB_ (Shrout and Fleiss, 1979). We used *r*_SB_ = 2*r*_split_*/*(1 + *r*_split_), where *r*_split_ is the correlation of participant-level scores estimated in the two trial halves. Confidence intervals were obtained by participant bootstrap resampling. Fixed- composition subsampling in the elevated-BDI group examined 8, 12, and 18 trials per valence. A descriptive higher-count series examined 20–100 trials per valence while allowing eligibility to change. The independent dataset used the same split definitions and fixed-count analyses at 20, 40, and 60 trials per valence. Split-half estimates quantify within-session consistency. They do not assess longitudinal test–retest reliability.

Electrophysiological characterization tested how narrowly the symptom association matched the FCz theta definition. Spatial analyses compared FCz, a frontocentral mean, a posterior control mean, and a 62-electrode topographic analysis (60 scalp electrodes and mastoids M1 and M2). Temporal estimates covered 0–200, 200–400, and 400–600 ms. Spectral analyses used delta (1–4 Hz), theta (4–8 Hz), alpha (8–12 Hz), and beta (13–20 Hz). P3-minus-N2 served as a time-domain event-related potential (ERP) comparator. Primary-dataset feature models used the same joint STAI-plus-BDI specification as the symptom analysis. Holm correction was used for the small families of planned contrasts (Holm, 1979). The 62-electrode analysis used the Benjamini–Hochberg false discovery rate (FDR) procedure because it involved simultaneous testing across electrodes (Benjamini and Hochberg, 1995). The independent dataset used the same spatial, temporal, spectral, and ERP definitions.

### 2.6 Computational identifiability and recovery

The computational analyses separated predictor overlap, population model discrimination, participant- level neural-slope recovery, and recovery of fitted reinforcement learning (RL) latent variables and parameters because group-level separability does not guarantee dependable individual-difference estimates (Wilson and Collins, 2019). We use practical computational identifiability to mean the ability of the available design and noisy observations to distinguish neural contributions of feedback valence and signed PE. Recovery instead measures how accurately known generating quantities are retrieved in simulation. Correct population-level model selection and accurate participant-specific slope recovery are distinct outcomes. Empirical valence–PE overlap was summarized using cor- relations and variance inflation factors (VIFs) after removing participant or session means when fixed effects made within-person geometry the relevant design. For a two-predictor diagnostic, VIF = 1*/*(1 *− r*^2^), where *r* is the relevant predictor correlation after accounting for the fixed-effects structure.

Exact-design simulations preserved the primary 46-participant, 11,025-trial design and generated neural outcomes from feedback valence, signed PE, joint, or null mechanisms across three observation- noise levels. Candidate estimators included valence-only, PE-only, and joint models. Population model discrimination, quantified by the correct model-selection rate, and participant-level neural- slope recovery were evaluated separately. A broader simulation grid varied the valence–PE correlation, balanced trial count, estimated-PE fidelity, and observation noise. Target correlations ranged from *−*0.965 to 0.965, trial counts ranged from 20 to 100 per valence, and estimated-PE fidelity, which is the within-participant correlation of simulated true and estimated PE regressors, was 1.0, 0.8, or 0.5. Each grid cell used 750 replications with 46 simulated participants. For single-source valence and signed-PE generators, “stable identification” was a simulation-specific four-part criterion: a mean across-participant neural-slope recovery correlation of at least 0.60, detection of the target STAI association in at least 60% of replications, an estimable target coefficient for at least 95% of simulated participants, and detection of the competing adjusted predictor in no more than 20% of replications. Unlike the separate population model-discrimination analysis, this grid criterion did not require correct model selection. These criteria summarize the simulation grid and are not empirical quality thresholds. Full simulation definitions are provided in Supplementary Methods.

Participant-wide RL latent-variable and parameter recovery used all 122 primary-dataset participants. Synthetic choices were generated from each participant’s fitted separate-rate model and refit with the same estimation procedure. Recovery was assessed for signed PE, within-valence signed PE, chosen value, positive-feedback PE, learning rates, inverse temperature, and learning-rate asymmetry.

For the independent dataset, predictor geometry was evaluated under one-rate and dual-rate learning models. Separate exact-design simulations assessed population model discrimination, measured by correct selection of the generating model, and recovery of participant-specific neural slopes.

### 2.7 Independent reproduction and model comparison

The independent reproduction analysis estimated the negative-minus-positive FCz theta contrast separately in Study 1 and placebo using participant-session fixed effects. Repeated trials within a participant-session violate independence at the trial level, so inference used the CR1 finite-sample correction to cluster-robust covariance estimates (Cameron and Miller, 2015). We report CR1 standard errors (SEs) and 95% confidence intervals (CIs), together with a two-sided sign-flip test and participant-session bootstrap CIs.

External neural encoding models contained feedback valence alone, signed PE alone, or both predictors. Incremental variance was summarized by partial *R*^2^. Predictive comparison used leave-one-session-out cross-validation, which was also leave-one-participant-out in the analyzed data. Population slopes were estimated on the remaining sessions and evaluated on within-session theta variation in the held-out session. The predictive comparison respected the session-level clustering of trials. Ordinary trial-level likelihood criteria do not represent this within-session dependence and were retained only as descriptive sensitivity measures. Paired held-out log-likelihood differences were summarized using session bootstrap intervals and sign-flip tests. The Akaike information criterion (AIC) and Bayesian information criterion (BIC) are reported as descriptive sensitivity measures in the supplement.

STAI was unavailable in the independent dataset. A secondary Study 1 analysis related BIS to theta gain while controlling BDI using partial Spearman correlation, permutation inference, and bootstrap CIs. BIS was treated as a related but non-equivalent construct.

All trial-level regressions used participant or participant-session fixed effects and CR1 cluster- robust covariance estimates (Cameron and Miller, 2015). The STAI-by-valence interaction was also evaluated with a restricted wild-cluster bootstrap because the symptom analysis involved a modest number of participant clusters (Cameron et al., 2008). Tests were two-sided. Holm correction was applied within prespecified families of related tests (Holm, 1979).

## 3 Results

### 3.1 Trait anxiety and feedback valence

The conventional signed-PE benchmark reproduced the previously reported symptom association in the 46 participants with elevated BDI scores (11,025 trials). Higher STAI was associated with more negative signed-PE coupling (*β* = *−*0.132, CR1 SE = 0.041, 95% CI [*−*0.213*, −*0.050], *p* = 0.0023; Table 1). Because signed PE and feedback valence were strongly coupled, the principal analysis represented the observed feedback contrast directly and entered STAI and BDI in the same model.

**Table 1:**
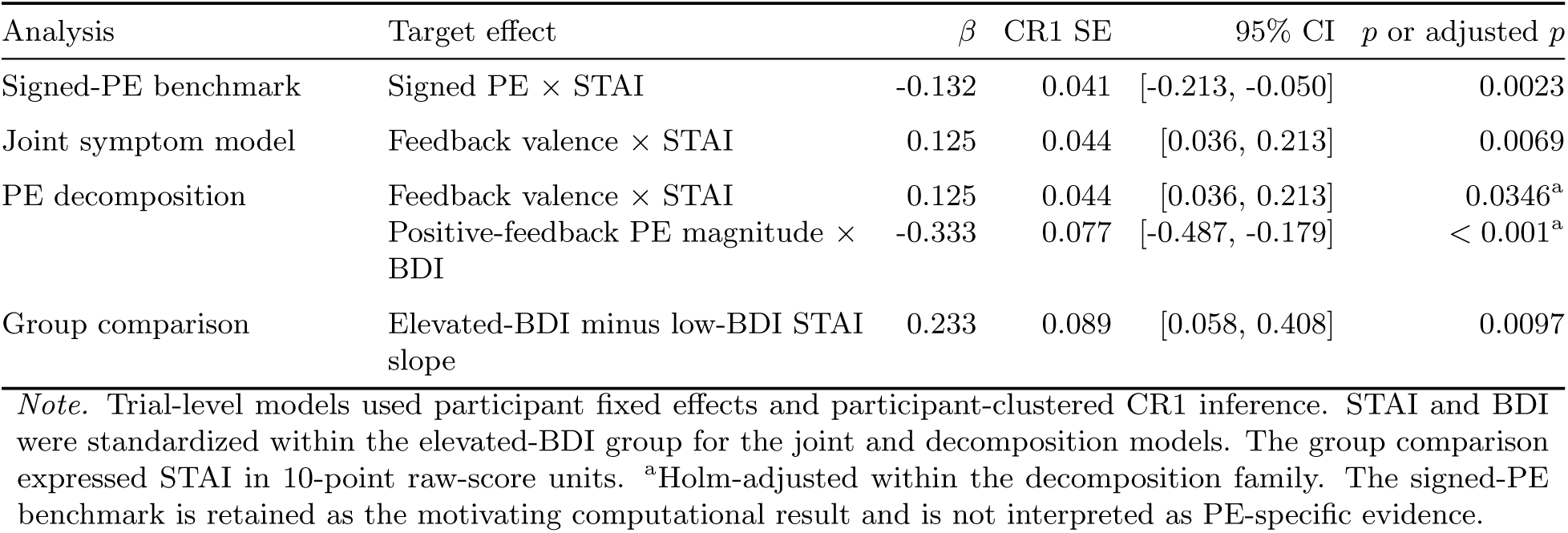
Symptom and computational-decomposition estimates in the primary dataset.

| Analysis | Target effect | *β* | CR1 SE | 95% CI | *p* or adjusted *p* |
|---|---|---|---|---|---|
| Signed-PE benchmark | Signed PE × STAI | -0.132 | 0.041 | [-0.213, -0.050] | 0.0023 |
| Joint symptom model | Feedback valence × STAI | 0.125 | 0.044 | [0.036, 0.213] | 0.0069 |
| PE decomposition | Feedback valence × STAI | 0.125 | 0.044 | [0.036, 0.213] | 0.0346<sup>a</sup> |
|  | Positive-feedback PE magnitude × BDI | -0.333 | 0.077 | [-0.487, -0.179] | < 0.001<sup>a</sup> |
| Group comparison | Elevated-BDI minus low-BDI STAI slope | 0.233 | 0.089 | [0.058, 0.408] | 0.0097 |

*Note.* Trial-level models used participant fixed effects and participant-clustered CR1 inference. STAI and BDI were standardized within the elevated-BDI group for the joint and decomposition models. The group comparison expressed STAI in 10-point raw-score units. <sup>a</sup>Holm-adjusted within the decomposition family. The signed-PE benchmark is retained as the motivating computational result and is not interpreted as PE-specific evidence.

STAI and BDI were moderately correlated within the elevated-BDI group (*r* = 0.399, *p* = 0.0060; VIF = 1.19). The STAI-by-valence coefficient was positive after adjustment for BDI (*β* = 0.125, CR1 SE = 0.044, 95% CI [0.036, 0.213], *p* = 0.0069), while the BDI-by-valence coefficient was smaller (*β* = 0.054, *p* = 0.209; Figure 1A). Restricted wild-cluster inference also supported the STAI interaction (*p* = 0.019). All 46 leave-one-participant-out estimates remained positive. The association was similar with raw-dB theta, an alternative baseline, and after removing response-time and trial-progression covariates (Supplementary Results).

**Figure 1:**
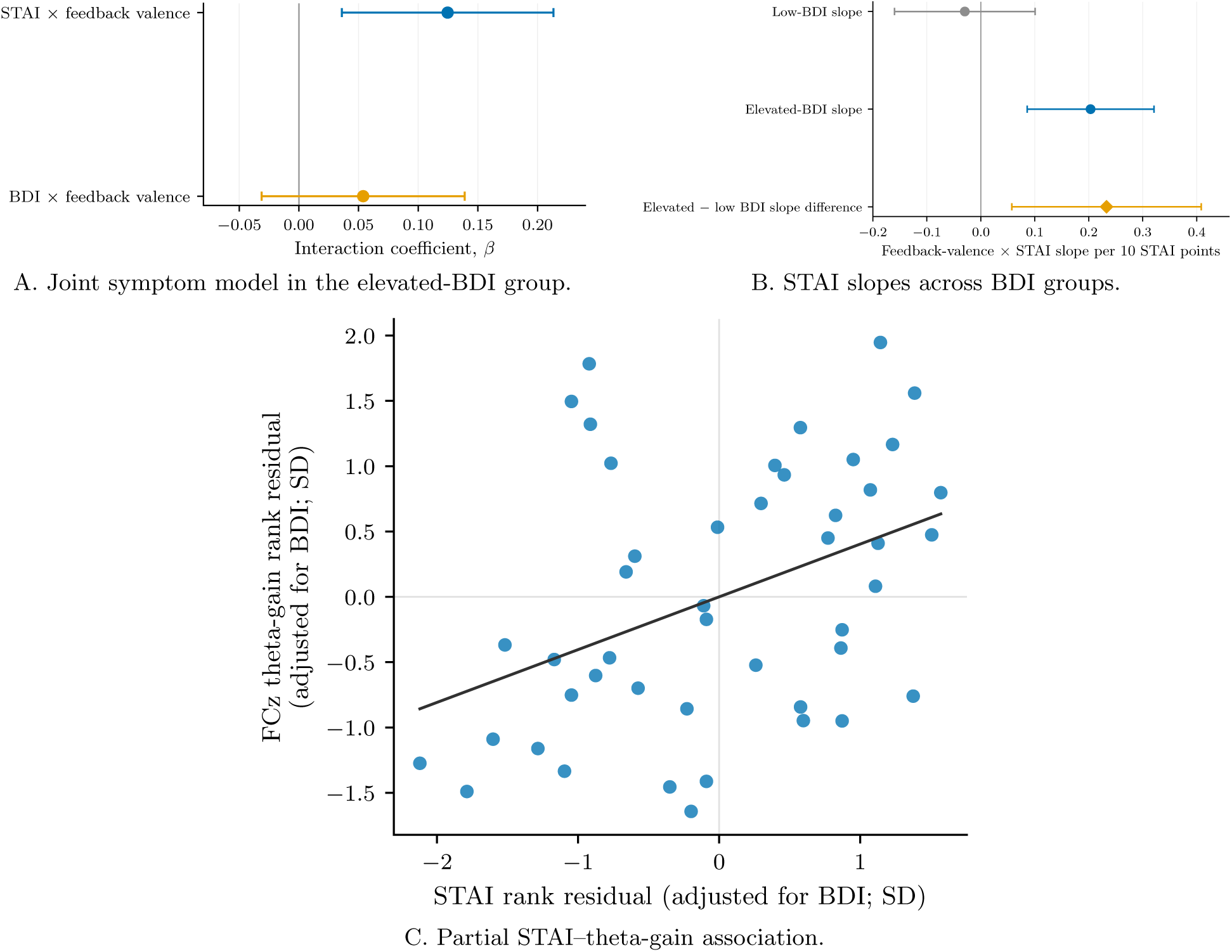
Trait anxiety and the feedback-valence response. (A) STAI-by-valence and BDI-by-valence coefficients from the joint model in the elevated-BDI group. (B) STAI-by-valence slopes in the low-BDI and elevated-BDI groups and their formal difference. Slopes are expressed per 10 STAI points. (C) Participant-level association between STAI and theta gain after adjustment for BDI. Error bars in panels A–B are CR1 95% confidence intervals. The line in panel C is descriptive.

Adding positive- and negative-feedback PE magnitude did not reduce the STAI-by-valence coefficient (*β* = 0.125, Holm-adjusted *p* = 0.0346). Neither STAI-by-PE-magnitude interaction was supported. BDI showed a separate association with positive-feedback PE magnitude (*β* = *−*0.333, CR1 SE = 0.077, 95% CI [*−*0.487*, −*0.179], Holm-adjusted *p <* 0.001). The equivalent chosen-value coefficient changed sign, as expected from *|δ*^+^*|* = 1 *− Q*, and remained supported under one-rate and separate-rate models, although its magnitude varied across models. The STAI-by-valence coefficient changed little across these specifications (Supplementary Figure S1). The coefficients are conditional associations within the elevated-BDI group and do not establish separate anxiety and depression mechanisms.

### 3.2 Symptom context and behavioral relevance

The low-BDI and elevated-BDI groups differed in the association between STAI and feedback valence. Expressing STAI in 10-point units, the feedback-valence slope was small in the low-BDI group (*β* = *−*0.030, SE = 0.066, *p* = 0.653) and positive in the elevated-BDI group (*β* = 0.203, SE = 0.059, *p <* 0.001; Figure 1B). The group-by-STAI-by-valence interaction was supported (*β* = 0.233, SE = 0.089, 95% CI [0.058, 0.408], *p* = 0.0097).

At the participant level, theta gain was associated with STAI after adjustment for BDI (partial Spearman *ρ* = 0.404, *p* = 0.0059; Figure 1C). The corresponding raw-dB association was similar. STAI residualized on BDI within the elevated-BDI group also retained a positive trial-level valence association. Larger theta gain was associated with a larger NoGo-minus-Go learning bias (*ρ* = 0.379, Holm-adjusted *p* = 0.0294). The separate Go and NoGo associations were less precise. The behavioral association is shown in Supplementary Figure S3 as secondary evidence of behavioral relevance. No mediation inference is made.

### 3.3 Reliability and electrophysiological characteristics

FCz theta gain showed useful within-session consistency in the elevated-BDI group. Spearman– Brown reliability was *r*_SB_ = 0.755 for odd–even splitting and 0.645 for first–second splitting (Table 2).

**Table 2:**
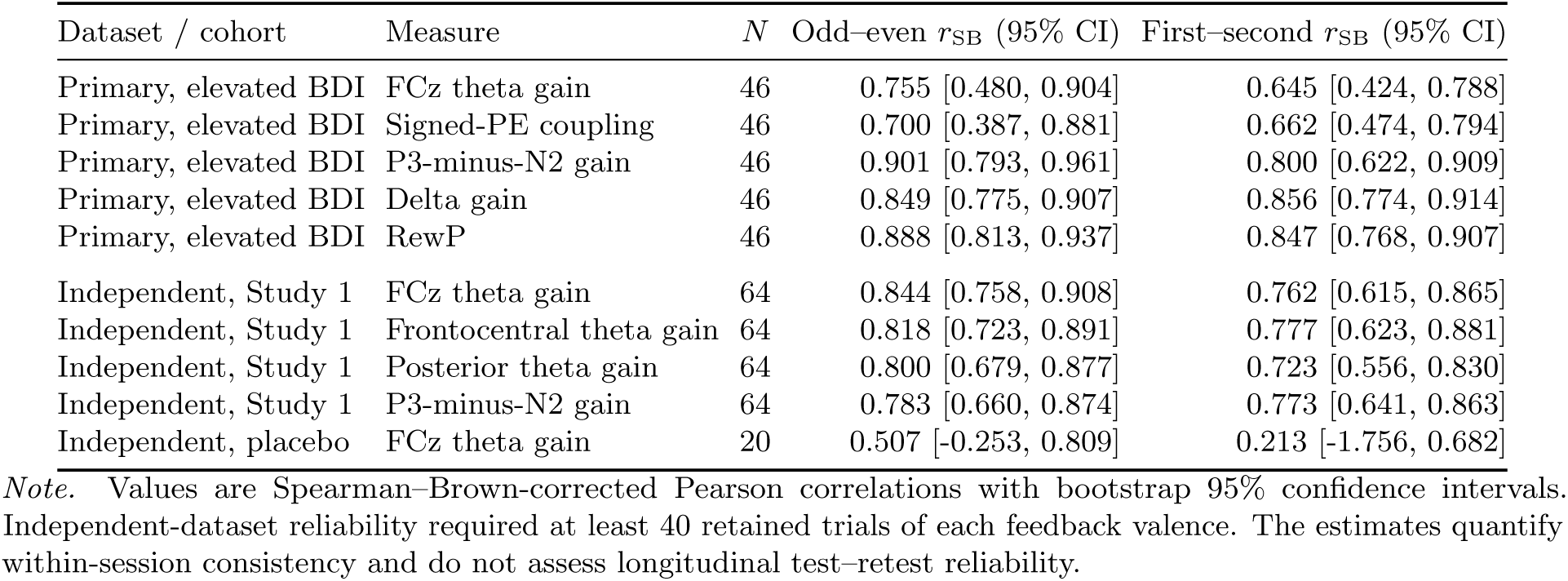
Within-session split-half reliability of feedback-related measures.

| Dataset / cohort | Measure | *N* | Odd–even *r*<sub>SB</sub> (95% CI) | First–second *r*<sub>SB</sub> (95% CI) |
|---|---|---|---|---|
| Primary, elevated BDI | FCz theta gain | 46 | 0.755 [0.480, 0.904] | 0.645 [0.424, 0.788] |
| Primary, elevated BDI | Signed-PE coupling | 46 | 0.700 [0.387, 0.881] | 0.662 [0.474, 0.794] |
| Primary, elevated BDI | P3-minus-N2 gain | 46 | 0.901 [0.793, 0.961] | 0.800 [0.622, 0.909] |
| Primary, elevated BDI | Delta gain | 46 | 0.849 [0.775, 0.907] | 0.856 [0.774, 0.914] |
| Primary, elevated BDI | RewP | 46 | 0.888 [0.813, 0.937] | 0.847 [0.768, 0.907] |
| Independent, Study 1 | FCz theta gain | 64 | 0.844 [0.758, 0.908] | 0.762 [0.615, 0.865] |
| Independent, Study 1 | Frontocentral theta gain | 64 | 0.818 [0.723, 0.891] | 0.777 [0.623, 0.881] |
| Independent, Study 1 | Posterior theta gain | 64 | 0.800 [0.679, 0.877] | 0.723 [0.556, 0.830] |
| Independent, Study 1 | P3-minus-N2 gain | 64 | 0.783 [0.660, 0.874] | 0.773 [0.641, 0.863] |
| Independent, placebo | FCz theta gain | 20 | 0.507 [-0.253, 0.809] | 0.213 [-1.756, 0.682] |

*Note.* Values are Spearman–Brown-corrected Pearson correlations with bootstrap 95% confidence intervals. Independent-dataset reliability required at least 40 retained trials of each feedback valence. The estimates quantify within-session consistency and do not assess longitudinal test–retest reliability.

Signed-PE coupling was similar, while P3-minus-N2, delta, and RewP were at least as reliable under one or both split schemes. Reliability declined when fewer trials were retained. With the 46-participant sample fixed, median reliability increased from 0.239 at eight trials per valence to 0.470 at 18 trials per valence. In the higher-count series, reliability increased further as the eligible sample became smaller (Figure 2A).

**Figure 2:**
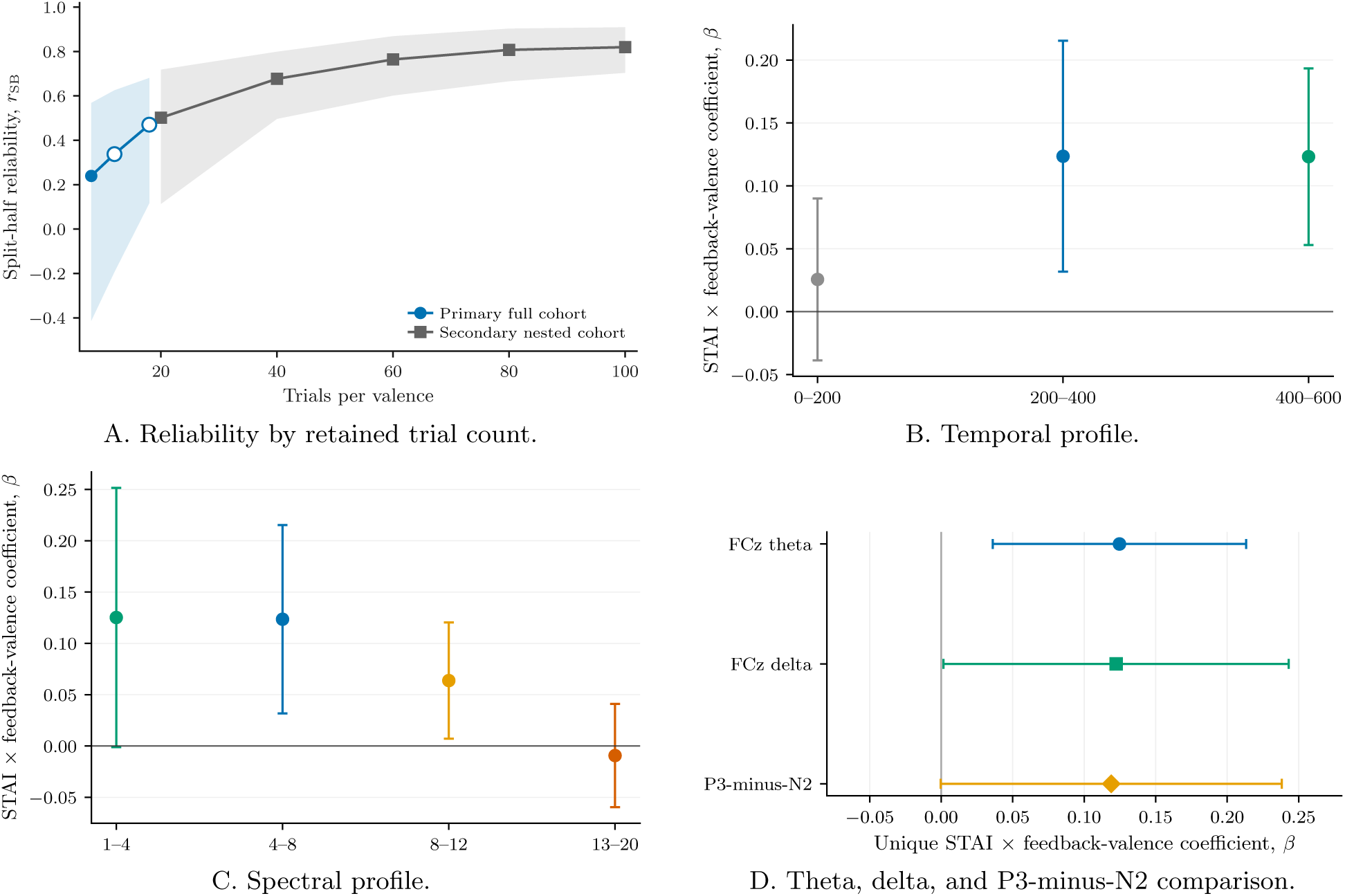
Reliability and electrophysiological characteristics of the primary response. (A) Median Spearman–Brown reliability across trial-count subsampling analyses. The higher-count series changes both trial count and sample composition. (B–C) STAI-by-valence coefficients across temporal windows and frequency bands. (D) Matched coefficients for theta, delta, and P3-minus-N2. Error bars are CR1 95% confidence intervals.

The symptom association was not sharply localized in space, time, or frequency. The STAI-by- valence coefficient was approximately 0.125 at FCz and 0.124 for the frontocentral average. The posterior estimate was smaller, but the frontocentral-minus-posterior contrast was unsupported after correction. No individual channel survived FDR correction. Participant-deletion and interpolation checks did not identify a single-subject or interpolation explanation for the descriptive frontocentral pattern (Supplementary Figure S2).

Temporally, the coefficient was small from 0–200 ms and similar in the 200–400- and 400–600-ms intervals (Figure 2B). Spectrally, delta and theta estimates were nearly identical, with smaller alpha and near-zero beta estimates (Figure 2C). No theta-minus-band contrast survived correction. P3-minus-N2 produced a similar STAI-by-valence coefficient, and direct theta-minus-delta and theta-minus-P3-minus-N2 contrasts were unsupported (Figure 2D). The prespecified FCz theta measure captures the symptom association, but the association is not confined to theta.

### 3.4 Computational identifiability and latent recovery

Next, we examined whether the primary task could distinguish neural contributions of feedback valence and signed PE. After removal of participant means, their correlation was approximately *r* = *−*0.964, with VIF = 14.06 (Supplementary Figure S13). In the exact-design simulations, single-predictor models readily assigned neural signal to the correlated competing predictor. Joint models improved population-level attribution, but recovery of unique participant-level neural slopes remained weak (Figure 3A). Population model selection could still identify the generating model in more than 99% of the same simulations. These findings are compatible: population selection aggregates evidence across trials and participants, whereas neural-slope recovery asks whether the individual generating coefficients can be estimated accurately.

**Figure 3:**
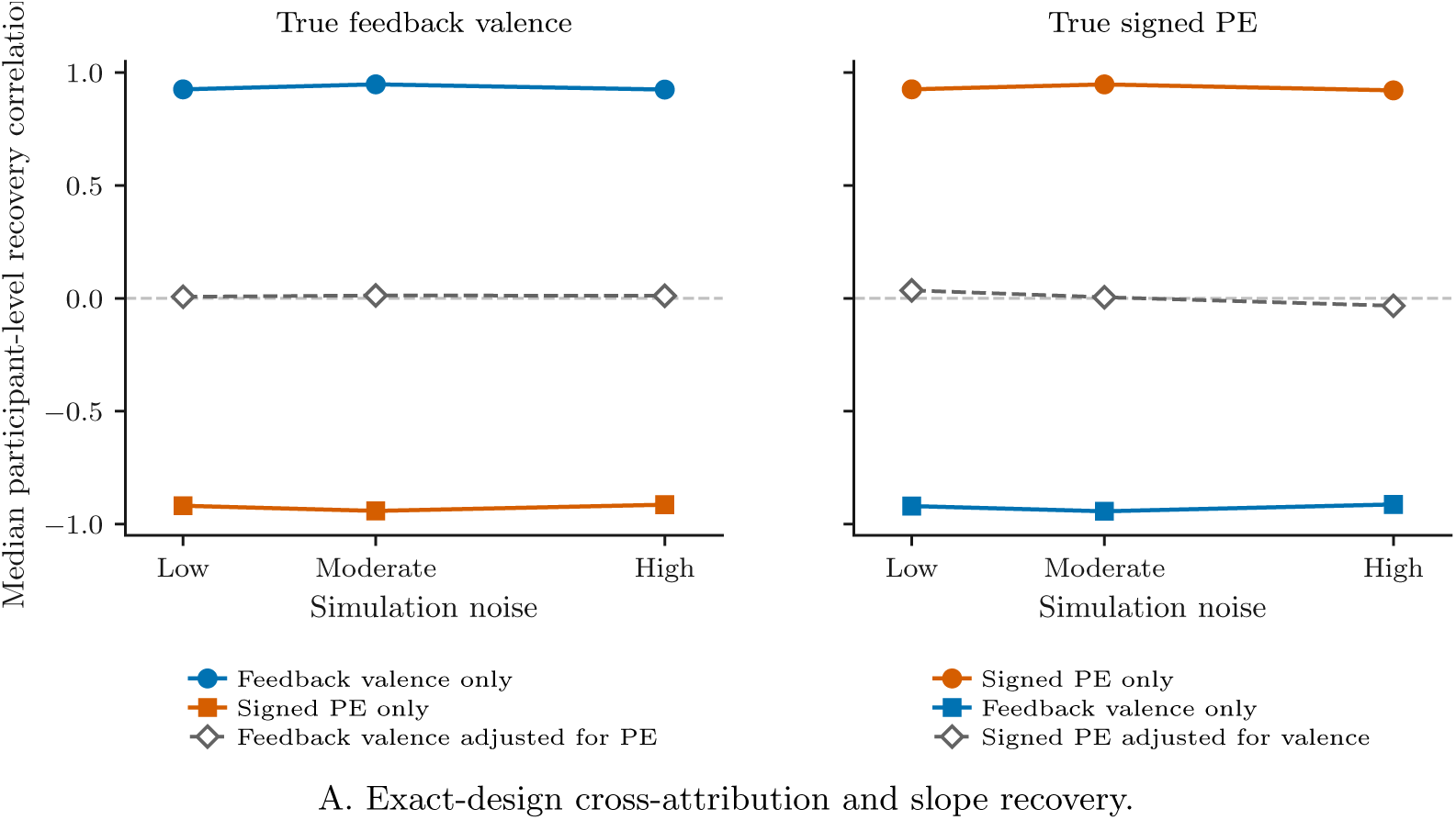
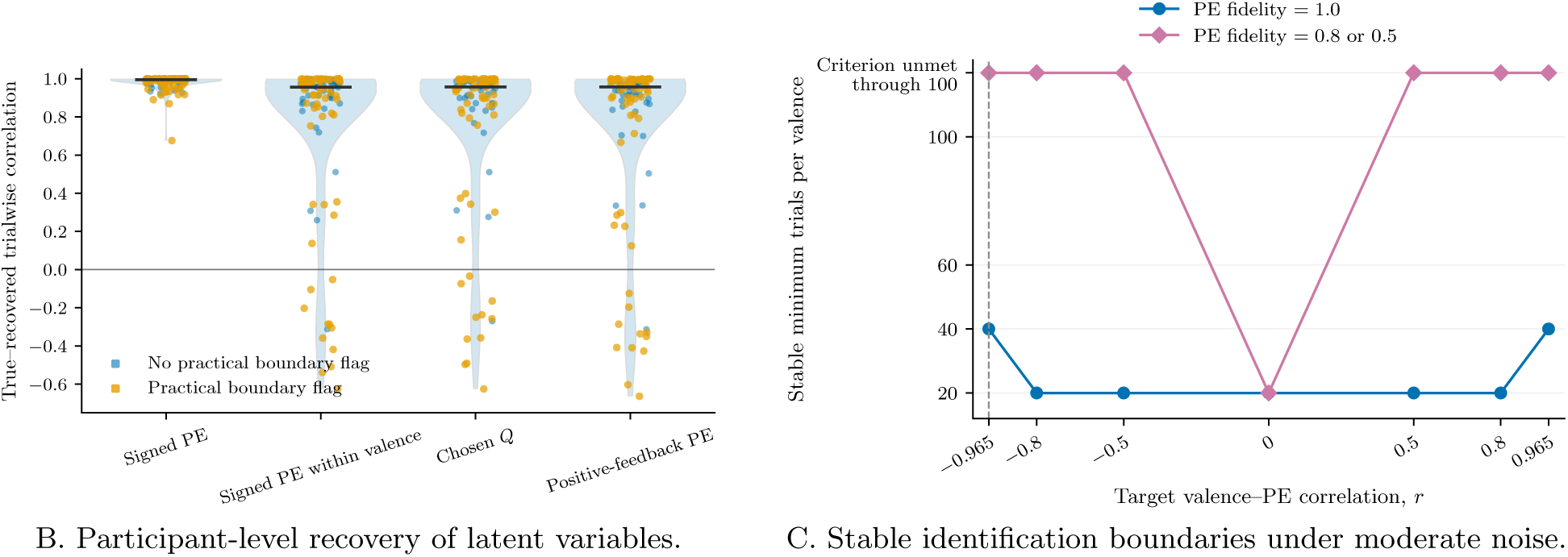
Computational identifiability and recovery in the primary task. (A) Correlations of estimated participant-level coefficients with known generating neural slopes under true-valence and true-signed-PE simulations. Fits using the competing predictor illustrate cross-attribution rather than recovery of a true competing effect. Panels B–C continue with the participant-level and design-boundary analyses. (B) True-to-recovered correlations for fitted RL latent variables after simulated-choice generation and model refitting across participants. (C) Minimum tested trial counts meeting the simulation-defined four-part signed-PE identification criterion. Estimated-PE fidelity denotes the correlation between simulated true and estimated PE regressors. Fidelities 0.8 and 0.5 had the same boundary. The upper category indicates that the criterion was not met through 100 trials per valence. The empirical valence–PE distribution is shown in Supplementary Figure S13.

Participant-wide RL latent-variable recovery provided a separate test of the fitted reinforcement- learning variables (Figure 3B). Overall signed PE showed high true-to-refit correlation, but this measure inherits information from the observed feedback sign. Within-valence signed PE, chosen value, and positive-feedback PE showed more heterogeneous recovery, including a lower tail with poor or negative true-to-recovered correlations. Learning-rate recovery was moderate, while inverse temperature and learning-rate asymmetry were less dependable. The fitted latent variables were adequate for the population analyses, but participant-specific estimates were not uniformly precise.

The broader simulation grid showed that signed-PE criterion attainment depended on predictor overlap, estimated-PE fidelity, trial count, and observation noise. Target-coefficient estimability was universal in the tested grid. Detection of the competing adjusted predictor was the most restrictive component of the four-part, simulation-defined criterion. At *|r|* = 0.965 and estimated-PE fidelity of 1.0, stable identification was achieved at 20, 40, and 100 trials per valence under low, moderate, and high noise, respectively. With estimated-PE fidelity of 0.8 or 0.5, no tested count through 100 trials per valence met the stable-identification criterion at the empirical-like overlap. Figure 3C shows the moderate-noise boundary. These numerical boundaries apply only to the simulated grid.

### 3.5 Independent reproduction of the feedback response

The negative-minus-positive FCz theta response reproduced in both independent cohorts (Figure 4A). Study 1 included 70 untreated sessions and 15,747 retained trials. The fixed-effects estimate was *β* = 0.364 (CR1 SE = 0.038, 95% CI [0.288, 0.440], *p <* 0.001), and 65 of 70 session contrasts were positive. Placebo included 23 sessions and 3,565 trials. Its estimate was *β* = 0.297 (CR1 SE = 0.050, 95% CI [0.193, 0.401], *p <* 0.001), and all 23 session contrasts were positive. Sign-flip tests supported both distributions. These analyses reproduce the neural feedback contrast, not the STAI association, which could not be tested because STAI was unavailable.

**Figure 4:**
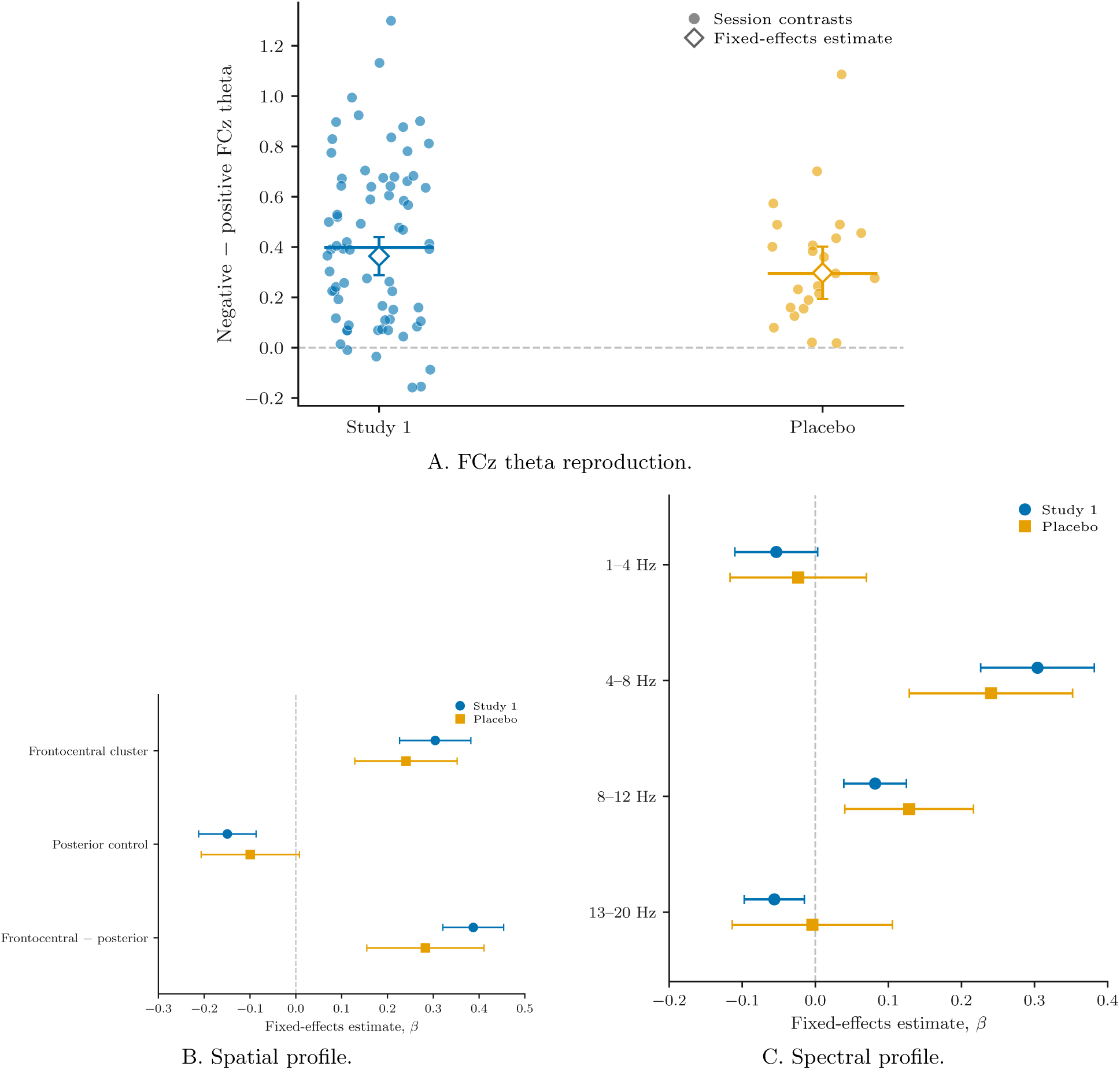
Independent reproduction and electrophysiological characteristics. (A) Participant-session negative-minus-positive FCz theta contrasts and fixed-effects estimates. (B) Frontocentral, posterior, and frontocentral-minus-posterior estimates. (C) Delta, theta, alpha, and beta estimates in the frontocentral 200–400-ms window. Error bars are CR1 95% confidence intervals.

Study 1 also showed strong within-session reliability (Table 2). Placebo reliability estimates were less precise because of the smaller eligible sample. The independent spatial profile was frontocentral in both cohorts (Figure 4B). Theta was more prominent than neighboring frequency bands in Study 1 and exceeded delta and beta in placebo, although not alpha (Figure 4C). The 200–400-ms interval was strongest in Study 1, and P3-minus-N2 showed the same feedback direction in both cohorts (Supplementary Results). The fixed neural measure reproduced across distinct recordings, whereas its spectral profile differed between datasets.

### 3.6 External computational model comparison

The independent task reduced valence–PE overlap under the one-rate Study 1 model but did not remove it. The within-session correlation was *r* = *−*0.889 with VIF = 4.78. Under the dual-rate model, overlap increased to *r* = *−*0.947 with VIF = 9.72. Placebo correlations were approximately *−*0.95 under both models (Figure 5A). Exact-design simulations under the one-rate Study 1 geometry showed near-perfect population model discrimination, whereas participant-specific neural slopes were recovered only moderately (Figure 5C).

**Figure 5:**
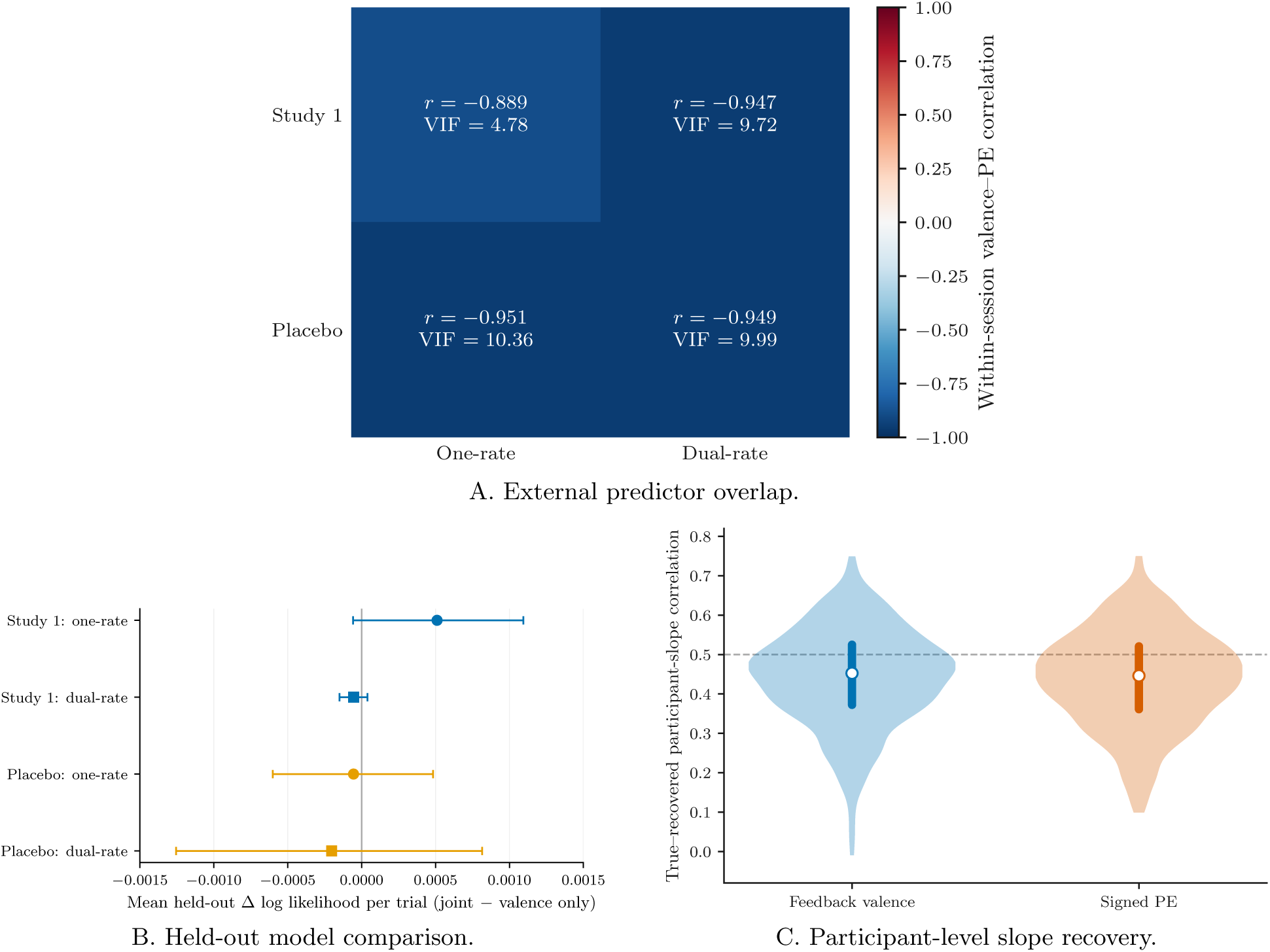
External computational analyses. (A) Valence–signed-PE overlap under one-rate and dual-rate learning models. (B) Mean held-out log-likelihood difference per trial for the joint minus valence-only model. Positive values favor the joint model. (C) Correlations between true and recovered participant-specific neural slopes in simulation.

In Study 1, the one-rate joint model contained a positive valence coefficient and a smaller signed-PE coefficient. Signed PE explained little additional variance after valence (partial *R*^2^ = 0.00118). Leave-one-session-out prediction showed a small aggregate gain for the joint model over valence alone, but the session-bootstrap interval included zero and the paired sign-flip test was unsupported (Figure 5B; Table 3). No comparable held-out predictive improvement was observed under the dual-rate Study 1 model or in placebo under either learning model. PE-only models did not outperform valence-only models in any external comparison. Trial-level information criteria showed the same dependence on learning-model specification and are reported as a descriptive sensitivity analysis in Supplementary Table S1.

**Table 3:** External joint-model coefficients, incremental variance, and prediction.

| Cohort | PE model | Valence β | PE β | R<sup>2</sup><sub>PE\|V</sub> | R<sup>2</sup><sub>V\|PE</sub> | Held-out Δ log L | Sign-flip p |
|---|---|---|---|---|---|---|---|
| Study 1 | One-rate | 0.226 | -0.076 | 0.001178 | 0.002527 | 8.017 | 0.0873 |
| Study 1 | Dual-rate | 0.334 | -0.015 | 0.000025 | 0.002852 | -1.337 | 0.2766 |
| Placebo | One-rate | 0.426 | 0.066 | 0.000420 | 0.004158 | -0.442 | 0.8478 |
| Placebo | Dual-rate | 0.472 | 0.090 | 0.000817 | 0.005351 | -0.663 | 0.7203 |

*Note.* Coefficients are from joint participant-session fixed-effects models. Partial R<sup>2</sup><sub>PE|V</sub> is the incremental variance explained by signed PE after feedback valence; R<sup>2</sup><sub>V|PE</sub> is the converse. Held-out Δ log L is joint minus valence-only log likelihood summed across leave-one-session-out folds. Positive values favor the joint model.

In a separate related-trait analysis, BIS was positively associated with theta gain after adjustment for BDI in Study 1 (partial Spearman *ρ* = 0.283, *p* = 0.0218). BIS is not equivalent to STAI, so this association is reported as related-trait evidence and not as replication of the primary symptom result (Supplementary Figure S12).

## 4 Discussion

We re-examined an association between anxiety and feedback-related EEG to assess the support for its interpretation in terms of fitted prediction error. In the elevated-BDI group, the STAI-by-valence association persisted after BDI adjustment and differed from the association in the low-BDI group. The prespecified FCz feedback contrast showed useful within-session consistency and reproduced in two independent cohorts, although STAI was unavailable for a direct symptom replication. Computational attribution was less secure. Valence and PE were closely coupled in the primary task, recovery of the within-valence latent information was heterogeneous, and the small external PE increment depended on the learning model.

### 4.1 Symptom and computational interpretation

The signed-PE analysis reproduced the symptom association reported by Cavanagh et al. (2019), but signed PE and feedback valence were nearly collinear within participants. When the direct feedback contrast and within-valence PE magnitude were modeled together, STAI remained associated with feedback valence and showed little evidence of association with PE magnitude. This finding does not imply that PE is absent from feedback-related EEG. In the exact-design simulations, a response generated by signed PE was also captured by a valence-only model, while the PE effect conditional on valence was recovered poorly. The empirical decomposition cannot determine which of these correlated quantities generated the neural response. Prediction-error accounts remain prominent in the feedback literature (Cavanagh et al., 2010; Holroyd and Coles, 2002; Sambrook and Goslin, 2015), and outcome valence, expectancy, and statistical context can contribute to overlapping neural responses (Hoy et al., 2021; Nassar et al., 2019; Stolz et al., 2023; Watts et al., 2017). Within the primary task, symptom variation was evident in the direct feedback contrast, while variation in PE magnitude within feedback valence contributed little to the STAI association.

The association with STAI occurred in a sample selected for elevated depressive symptoms. STAI predicted the feedback contrast after BDI adjustment within that group, and the STAI-by-valence slope was larger there than in the low-BDI group. Participant-level analyses showed a similar BDI-adjusted association. The result places anxiety-related variation in an elevated-depressive-symptom context and does not establish an anxiety-specific biomarker independent of depression. Katz et al. (2020) found substantial overlap between depression and anxiety in reinforcement sensitivity, and computational studies across both symptom domains remain heterogeneous (Pike and Robinson, 2022). EEG studies that measure both symptom domains likewise report nonuniform relationships with feedback processing (Grabowska et al., 2024; Rohlinger et al., 2026).

BDI was associated with the positive-feedback latent variable after STAI adjustment. The coefficient reversed sign under the chosen-value parameterization, as expected from its inverse relationship with positive-feedback PE magnitude. Its magnitude varied across learning models, and the corresponding associations were not supported for the tested delta or P3-minus-N2 outcomes. We treat this as a secondary computational observation. The data do not support a separate depression mechanism.

### 4.2 Measurement properties and external generalization

The direct feedback response was measurable at the participant level. Split-half reliability in the primary elevated-BDI group was adequate for cautious individual-difference analysis and improved with greater trial availability. Other feedback-related measures were at least as reliable under some split schemes, so reliability alone does not privilege FCz theta. This dependence on score construction and measurement design is consistent with psychometric work on ERP and reward-related difference measures (Amir et al., 2023; Clayson and Miller, 2017; Meyer et al., 2017). The estimates reported here address within-session consistency and not longitudinal stability. These reliability estimates apply to the FCz feedback contrast. Frequency specificity and recovery of a fitted participant-level EEG–PE slope require separate evidence.

The electrophysiological characterization also limits a narrow theta interpretation. Delta and theta produced similar symptom coefficients, P3-minus-N2 showed a comparable effect, and no individual channel survived FDR correction in the primary symptom analysis. The temporal response extended across 200–600 ms. FCz theta remains a useful prespecified measurement target, but the symptom association appears to be part of a broader feedback-related response. Time-domain and low- frequency feedback measures are known to overlap and to vary with task context (Bernat et al., 2015; Hoy et al., 2021; Williams et al., 2021).

The independent dataset provides stronger evidence for the neural measurement than for its symptom interpretation. The negative-minus-positive FCz theta response reproduced in untreated Study 1 and placebo, and Study 1 showed strong split-half reliability and a clear frontocentral profile. The external response was more theta-prominent than the primary symptom effect. Differences in task structure, sample composition, acquisition, and artifact handling could each contribute to that spectral difference. Artifact procedures differed because the reference and auxiliary-channel configurations differed across datasets. The FCz channel, 4–8-Hz band, 200–400-ms latency, baseline interval, and dB normalization were kept constant. This design supports comparison of the prespecified neural measure, but preprocessing remains one possible contributor to cross-dataset spectral differences. STAI was unavailable in the independent dataset, so the anxiety association could not be replicated directly. The BIS analysis supplies related-trait information and does not substitute for an STAI replication. The primary NoGo-minus-Go association and the independent Study 1 BIS association provide distinct, secondary links between the feedback contrast and avoidance- or inhibition-related measures, although they do not constitute replication of the same behavioral relationship.

### 4.3 Implications for interpreting computational EEG associations

Established psychometric and computational checks answer different questions in this dataset. Split-half correlations characterize the consistency of the measured feedback contrast, valence– PE correlations quantify the predictors’ shared variation, and recovery analyses assess whether the remaining information supports estimates for individual participants. These checks narrow the computational interpretation of the previously reported symptom–EEG association without overturning the association itself. In the primary task, single-predictor models readily assigned simulated signal to the correlated competing predictor. Joint adjustment reduced attribution to the correlated competing predictor, but individual neural-slope recovery remained weak. In a separate analysis, population-level model selection identified the generating model in more than 99% of exact-design simulations. This did not establish accurate individual effect estimates. In the broader simulations, the four-part stable-identification criterion was met despite high predictor overlap under some tested conditions with perfect estimated-PE fidelity and sufficient trials. At empirical-like overlap, the criterion was not met at any tested count through 100 trials per valence when estimated-PE fidelity was 0.8 or 0.5.

Participant-wide reinforcement-learning latent-variable recovery clarifies what the full signed-PE correlation measures. Overall signed PE was recovered extremely well because its sign partly reflects observed binary feedback. Within-valence PE and chosen value were less dependable for a subset of participants. Near-ceiling recovery of the full signed regressor can coexist with weaker recovery of the latent variation needed to separate PE from feedback valence.

In the independent task, the one-rate Study 1 specification reduced predictor overlap and supported near-perfect population model discrimination in simulation, although participant-level neural-slope recovery remained moderate. Empirically, adding signed PE to valence yielded a small aggregate held-out gain under the one-rate Study 1 model, with limited consistency across sessions. No comparable gain was supported under the dual-rate model or in placebo. Predictor geometry is shaped by both the task and the latent-model specification. The observed PE increment is accordingly model dependent and does not provide unique evidence for a latent mechanism.

For studies using fitted learning variables to support participant-level symptom claims, we recommend assessing reliability of the neural score analyzed and, where feasible, recovery of the relevant participant-level coefficients. Additional trials can reduce sampling noise but do not necessarily supply the within-valence variation needed to distinguish PE from observed feedback. The trial-count boundaries in our simulation grid are conditional on the tested generating models, estimated-PE fidelity manipulation, and observation-noise assumptions.

## 5 Limitations and Future Directions

The primary symptom analysis included 46 participants with elevated BDI scores. The large number of trials improves estimation of within-participant neural effects but does not replace participant- level sample size for symptom inference. The source study also sampled separated low-BDI and elevated-BDI groups, so the group comparison does not estimate a continuous symptom function across the full BDI range.

The present work is a secondary analysis of open datasets, and severe predictor overlap in the primary task cannot be removed statistically. The independent task improved the geometry under one latent model but retained substantial collinearity, and the incremental signed-PE result changed with model specification. STAI was not collected in the independent dataset, preventing direct replication of the main symptom association. BIS measures behavioral inhibition and is not an equivalent anxiety measure.

Split-half reliability measures within-session consistency and does not establish stability across sessions or days. Artifact handling also differed between datasets because their recording references and auxiliary-channel configurations were different. A single artifact pipeline would not make the underlying acquisitions equivalent. We kept the neural target and time–frequency measurement definition constant and tailored artifact detection to each recording system. Cross-dataset spectral comparisons should still be interpreted cautiously because preprocessing may contribute to differences in spectral profile. All symptom and behavioral associations are observational and do not establish causal or clinical utility.

Future studies could use experimental designs that more independently manipulate feedback valence and expectation, while assessing how task structure, trial availability, and repeated measurements affect the reliability and recovery of participant-specific neural effects. Prospective simulations could also help evaluate whether a design provides sufficient information to distinguish competing computational explanations before data collection. More broadly, similar considerations may inform the use of machine-learning and reinforcement-learning models in psychophysiological research, where predictive performance, predictor distinctness, and recovery of individual-level quantities need not coincide. Applying these complementary checks may help distinguish useful predictive models from computational interpretations that the available physiological measurements cannot uniquely support.

## 6 Conclusion

Across two open EEG datasets, a negative-minus-positive frontocentral feedback response was reproducible and showed useful within-session consistency. Among participants with elevated depressive symptoms, trait anxiety was associated with the magnitude of this response after BDI adjustment. Matched low-frequency and time-domain analyses did not support theta specificity. The incremental information associated with signed PE was small and changed across learning models and independent cohorts. Feedback valence was the most reproducible empirical description supported by these data. The study does not establish a theta-specific mechanism, unique PE encoding, or a precise participant-level computational phenotype.

## Author Contributions

**Vaibhav Sinha:** conceptualization, data curation, formal analysis, investigation, methodology, writing - original draft, review & editing

**Vikram Ravindra:** conceptualization, supervision, writing - review & editing

## Ethics and Consent

The present study is a secondary analysis of publicly available, de-identified data and involved no new participant recruitment, intervention, or collection of identifiable human data. For the primary dataset, Cavanagh et al. (2019) report that all participants provided written informed consent approved by the University of Arizona. For the independent dataset, Cavanagh et al. (2014) report that both experiments were approved by the Brown University Institutional Review Board and that all participants provided written informed consent. No additional participant contact or data collection occurred for the present analyses.

## Data Accessibility

The primary EEG dataset is publicly available through OpenNeuro as ds003474, version 1.1.0 (Cavanagh, 2021). The independent EEG dataset is publicly available through OpenNeuro as ds004532, version 1.2.0 (Cavanagh and Frank, 2023).

