## Supplementary Material for "Feedback Valence, Prediction Error, and the Limits of Computational Attribution in Symptom-Related EEG"

#### 1 Supplementary Methods

##### 1.1 Additional EEG preprocessing details

For OpenNeuro dataset `ds003474`, electroencephalographic (EEG) processing retained 60 scalp channels, horizontal electrooculogram (HEOG) and vertical electrooculogram (VEOG) channels, and mastoid electrodes M1 and M2. The publicly released continuous EEGLAB files and event metadata were used as inputs to our own MNE-Python preprocessing and event-alignment procedures (Gramfort et al., 2013). The source study's MATLAB preprocessing code used the `APPLE_Dep` routine. That original routine was not used to create the present analysis epochs (Cavanagh et al., 2019). A standard 10–20 montage was assigned before interpolation. Continuous EEG and EOG were filtered at 0.5–100 Hz with a zero-phase finite-impulse-response filter (MNE-Python `firwin`) after a 60-Hz notch. Feedback-locked epochs extended from $-2$ to $+2$ s, without time-domain baseline subtraction. Bad-channel screening used log variance, the 95th percentile of epoch peak-to-peak amplitude, the 95th percentile of epoch maximum absolute amplitude, and negative correlation with the median scalp signal. Each metric was standardized across scalp channels within participant. A channel was flagged when any applicable, directionally defined metric had $z > 3$. Low correlation with the median scalp signal was assessed using the negative of that correlation. These were upper-tail screening rules, not $|z| > 3$ rules. Amplitude-based channel criteria were not applied to FP1, FPz, or FP2 before ocular-component removal. Log variance and correlation criteria remained active at those electrodes. More than six flagged scalp channels and interpolation of FCz were retained as participant-level quality-control indicators rather than automatic primary-dataset exclusions.

Epoch rejection used upper-tail $z > 3$ thresholds, standardized across epochs within participant, for mean absolute amplitude, log variance, maximum absolute amplitude, and maximum scalp-channel peak-to-peak amplitude. An epoch was rejected if any criterion was met. Independent component analysis (ICA) with FastICA (Hyvärinen and Oja, 2000) was fit to a 1-Hz high-pass copy of the retained epochs. ICA retained components accounting for 99% of variance (MNE-Python `n_components=0.99`, random state 97). Automated EOG component detection was performed separately for HEOG and VEOG at a threshold of 3.0. The union of identified components was excluded without manual component selection. A final signal-integrity rule removed epochs with median scalp-channel peak-to-peak amplitude below 5 $\mu$V in either the $-300$ to $-200$ ms baseline interval or the 200–400 ms measurement interval. This post-ICA check excluded eight epochs from one participant (27,311 before this check; 27,303 afterward). The rule was documented from EEG signal characteristics before symptom modeling.

For the primary dataset, stimulus markers defined trial windows ending before the next stimulus. Within each training window, the first feedback event and the last response preceding it were selected; later feedback markers were ignored. Following the audited event-label convention, right responses selected the first listed stimulus and left responses the second. Only trials with a selected choice, binary feedback, and valid stimulus–response–feedback timing entered the RL-eligible table. In the feedback-free test phase, the first response was used, and seven windows containing both left- and right-side responses were excluded from the strict behavioral table. The resulting canonical reconstruction contained 27,998 RL-eligible training trials before subsequent EEG screening. For EEG epoching, the reconstructed feedback onsets were converted to recording sample indices and linked to canonical trial metadata.

For `ds004532`, our pipeline read the public continuous EEGLAB `.set/.fdt` recordings (63 EEG channels and one EOG channel), assigned an MNE standard 10–05 montage, and applied a 60-Hz notch and zero-phase fourth-order Butterworth 0.5–40-Hz filtering for analysis. A 1–40-Hz copy was used for bad-channel detection after a 60-Hz notch. The acquisition used CPz as the online reference, which was not retained as a recorded channel, and the auxiliary EOG signal had different scaling from the primary release. Channels were flagged for nonfinite data, recording-relative robust-$z$ outliers ($|z| > 4$) in log median absolute deviation or log 0.5th–99.5th percentile range, or a near-flat signal on more than 5% of successive samples. No more than 10 EEG channels could be interpolated in a recording. Flagged channels were interpolated with spherical splines before average EEG rereferencing. A recording was excluded if FCz was flagged, more than 10 EEG channels were flagged, or the required montage and channel checks failed. FCz was not interpolated. Feedback epochs from $-0.8$ to $+1.0$ s were rejected when global EEG peak-to-peak amplitude exceeded 300 $\mu$V, FCz peak-to-peak amplitude exceeded 150 $\mu$V, or the recording-relative robust $z$ score of log EOG epoch peak-to-peak amplitude exceeded 4. No absolute EOG threshold was used. ICA was not applied because decomposition conditioning varied across recordings and did not provide a common artifact-removal rule for this release. The EEG-only pilot found markedly different principal-component variance concentrations across recordings, while EOG amplitude scaling also varied. The prespecified alternative used participant-relative EOG rejection and EEG/FCz peak-to-peak limits without ICA.

Artifact handling was tailored to the two acquisition systems. The primary release provided dedicated HEOG and VEOG channels and mastoid references that supported ICA-based ocular correction and mastoid rereferencing. The independent release had a different reference and auxiliary-channel configuration, so artifact control relied on recording-relative rejection criteria. Across datasets, the neural target remained FCz activity from 4–8 Hz and 200–400 ms, normalized in decibels to a pooled $-300$ to $-200$ ms baseline. The threshold choices were dataset-specific and documented before testing the symptom and neural-contrast associations. They are not presented as universal EEG cutoffs (Keil et al., 2014).

Time–frequency power in both datasets was estimated with complex Morlet wavelets on a 50-point logarithmic frequency grid from $10^{0.01}$ to $10^{1.7}$ Hz and a corresponding logarithmic cycle grid from three to 10 cycles. Wavelet convolution used the fast Fourier transform (FFT), no decimation, and the zero-mean convention. The eight grid frequencies between 4 and 8 Hz contributed to the theta measure. At these frequencies (4.27–7.45 Hz), the wavelets used approximately 4.67–5.55 cycles. For the Gaussian Morlet envelope, $\sigma_t = n_{\text{cycles}}/(2\pi f)$ was approximately 118–174 ms and $\sigma_f = 1/(2\pi\sigma_t)$ was approximately 0.92–1.34 Hz. Consequently, power estimates are smoothed around their nominal time points, including within the 100-ms baseline interval. The baseline window does not imply sharply bounded temporal support (Keil et al., 2022).

#### 1.2 Reinforcement learning estimation

Reinforcement-learning (RL) models used Q-learning action values $Q_t(a)$, where $t$ indexes trials and $a$ indexes an available action. Prediction error (PE) was the difference between the observed binary outcome and the pre-feedback chosen value. For the primary dataset, one-rate and separate-rate models were fit separately for each participant. Ten deterministic starts were evaluated per participant and model with Nelder–Mead optimization. Learning rates were represented through logistic transforms, and the inverse-temperature parameter used a positive softplus transform. Optimization allowed up to 10,000 iterations and 20,000 function evaluations with $10^{-8}$ parameter and objective tolerances. The best finite likelihood across starts was retained.

For the independent dataset, learning rates were bounded to $[0.001, 0.999]$, inverse temperature to $[0.01, 20]$, and $Q_0$ remained 0.5. The one-rate model used 12 deterministic starts spanning four learning rates and three inverse temperatures. The dual-rate model used 18 starts spanning three positive learning rates, three negative learning rates, and two inverse temperatures. Fits used the limited-memory Broyden–Fletcher–Goldfarb–Shanno algorithm with box constraints (L-BFGS-B), with a maximum of 2,000 iterations and objective tolerance $10^{-12}$.

#### 1.3 Additional symptom and behavioral analyses

The Beck Depression Inventory (BDI) and State–Trait Anxiety Inventory (STAI) were analyzed as described in the main article. We repeated the elevated-BDI symptom model in three sensitivity analyses: using theta in raw decibel (dB) units, using a within-trial baseline, and omitting response-time and trial-progression covariates. The within-trial baseline used each trial's frequency-specific mean power from $-300$ to $-200$ ms rather than the participant-pooled baseline for decibel normalization. Influence was assessed with leave-one-participant-out refits. A secondary analysis residualized STAI on BDI within the elevated-BDI group and substituted the residual score for STAI. This analysis asks whether within-group STAI variation not linearly associated with BDI remains related to the feedback contrast. It does not isolate a pure anxiety construct.

The positive-feedback latent association was evaluated in PE-magnitude and chosen-value coordinates under separate-rate and one-rate learning models. Since positive-feedback PE magnitude equals $1-Q$, these parameterizations carry the same trialwise information with opposite sign. P3-minus-N2 and delta outcomes were also tested to determine whether the latent association generalized beyond the principal theta measure.

For the behavioral analysis, participant-specific theta gain was correlated with NoGo-minus-Go, NoGo, and Go performance. Inference used 9,999 permutations and 9,999 participant bootstrap resamples. Holm correction was applied to the planned family. Bootstrap uncertainty is reported as 95% confidence intervals (CIs).

#### 1.4 Reliability procedures

Eligible EEG trials were ordered chronologically. Odd–even and first–second splits were then formed. When a session contained an odd number of trials, the final chronological trial was omitted from the split calculation only. Features were standardized over the full retained session before splitting. For direct theta gain, the participant-specific feedback-valence model was fit separately within each half. The signed-PE comparator replaced feedback valence with within-participant standardized signed PE. Response time and trial progression remained as covariates.

The primary statistic was the Spearman–Brown-corrected Pearson correlation. Raw Pearson and Spearman correlations, Spearman–Brown-corrected Spearman correlations, and intraclass correlation coefficients (ICCs), ICC(3,1) and ICC(3,2), were also recorded (Shrout and Fleiss, 1979). Bootstrap intervals used 9,999 participant resamples with common indices across measures. The primary fixed-composition trial-count series sampled 8, 12, or 18 trials per valence for all 46 elevated-BDI participants in 250 draws. A secondary nested series used 20, 40, 60, 80, and 100 trials per valence and allowed eligibility to decline with the required count. Independent-dataset fixed-count analyses used 20, 40, and 60 trials per valence with 250 draws.

#### 1.5 Simulation and recovery details

The exact-design primary simulations preserved the observed 46-participant, 11,025-trial design. Valence, signed-PE, mixed valence-plus-PE, and null generators were evaluated across low, moderate, and high observation-noise conditions. Secondary generators included unsigned surprise and a negative-feedback-gated absolute signed-PE term (zero on positive-feedback trials before standardization). For non-null single-source simulations, participant-specific signal slopes were generated as $0.35 + 0.275A_i + u_i$, where $A_i$ was standardized STAI and $u_i \sim N(0, 0.35^2)$. Observation-noise standard deviations were 0.75, 1.25, and 2.00. Each mechanism-by-noise cell used 120 replications.

The broader design grid varied target valence–PE correlation across −0.965, −0.80, −0.50, 0, 0.50, 0.80, and 0.965, balanced trial counts of 20, 40, 60, or 100 per feedback valence, estimated-PE fidelity of 1.0, 0.8, or 0.5, and three observation-noise levels. For each simulated participant, a standardized true signed-PE regressor $\delta^*_{it}$ was constructed with the prescribed correlation to feedback valence. An estimated PE regressor was then formed as

$$\begin{aligned} \widehat{\delta}^{(f)}_{it} &= f\,\delta^*_{it} + \sqrt{1-f^2}\,\eta_{it}, \\ \mathrm{corr}_t\left(\delta^*_{it}, \widehat{\delta}^{(f)}_{it}\right) &= f, \qquad f \in \{1.0, 0.8, 0.5\}, \end{aligned} \tag{S1}$$

where $\eta_{it}$ is a within-participant standardized random perturbation orthogonalized against the intercept, feedback valence, true PE, response time, and trial progression. This construction fixes the within-participant true–estimated PE correlation at $f$. Neural outcomes were generated from the true predictors. Ordinary estimators used the estimated PE, while oracle estimators used the true PE. Thus, fidelity represents controlled trialwise regressor misspecification, not the recovery rate of fitted RL parameters. Each cell used 750 replications with 46 simulated participants. For single-source valence or signed-PE generators, the simulation-specific “stable identification” criterion required (i) mean across-participant recovery correlation of at least 0.60 for known generating versus estimated neural slopes, (ii) detection of the target STAI association at $p < 0.05$ in at least 60% of replications, (iii) linear-algebra estimability of the target coefficient for at least 95%

of participants, and (iv) detection of the competing adjusted predictor in no more than 20% of replications. Model-selection accuracy was not part of this grid criterion. These thresholds were used to summarize the simulation grid, not as general EEG design standards.

Participant-wide RL latent-variable and parameter recovery generated synthetic choices from each fitted primary-dataset participant and refit the separate-rate model with the same multistart procedure. Recovery summaries included trialwise signed PE, signed PE within feedback valence, chosen value, positive-feedback PE, positive and negative learning rates, log inverse temperature, and learning-rate asymmetry.

For the independent dataset, population exact-design simulations used observed participant-session predictors with 1,000 replications per generating scenario. Separate participant-level simulations used 500 replications, participant-specific generating slopes with standard deviation 0.50, and unit Gaussian observation noise. Leave-one-session-out predictive comparisons demeaned outcome and predictors within each training session, estimated population slopes on the remaining sessions, and evaluated within-session theta variation in the held-out session.

### 2 Supplementary Results

#### 2.1 Symptom-model sensitivity analyses

The BDI-adjusted STAI-by-valence coefficient remained positive across the planned sensitivity analyses. With response-time and trial-progression covariates removed, the separate-rate coefficient was approximately 0.122 with $p = 0.012$ in the chosen-value model. In raw-dB units, the corresponding STAI-by-valence association remained positive and supported. Wild-cluster inference and leave-one-participant-out analyses preserved its direction. In the direct valence model without latent terms, all 46 leave-one-participant-out STAI coefficients remained positive.

Residualizing STAI on BDI within the elevated-BDI group also retained a positive STAI-by-valence association ($\beta = 0.097$, $p = 0.026$). This differs from residualization across the two separated BDI groups, which confounds the sampling structure with the symptom adjustment. The within-group result supports unique STAI-related variation after linear BDI adjustment without implying an anxiety-specific mechanism.

The BDI association with positive-feedback PE magnitude remained supported in a BDI-only decomposition. Under the chosen-value parameterization, its sign reversed as required by $|\delta^+| = 1-Q$. The association was present under one-rate and separate-rate models but was smaller under the one-rate specification. The STAI-by-valence coefficient was much less sensitive to the learning-model choice (Figure S1).

#### 2.2 Spatial characterization and behavioral associations

Figure S2 shows the 62-electrode STAI-by-valence map (60 scalp electrodes plus mastoids M1 and M2). Several frontocentral channels had positive descriptive coefficients, but no individual channel survived false discovery rate (FDR) correction. Participant-deletion fits retained positive coefficients for the largest frontocentral estimates, and preprocessing records did not indicate systematic interpolation at those channels.

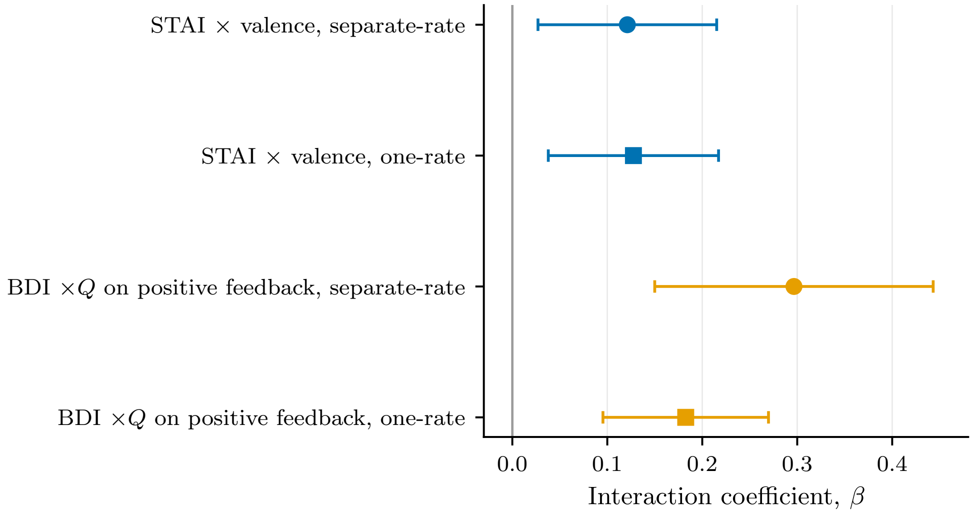


Figure S1: **Learning-model sensitivity of symptom associations.** STAI-by-valence and BDI-by-positive-feedback chosen-value coefficients under separate-rate and one-rate Q-learning models.

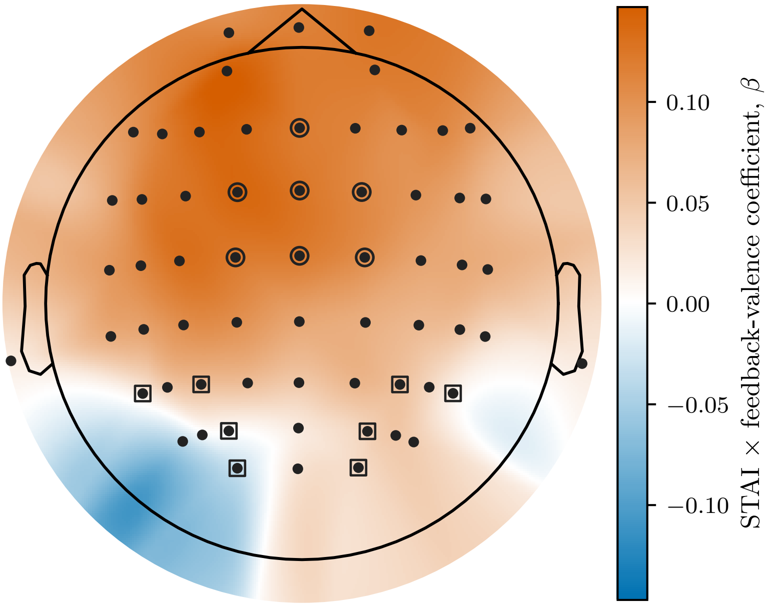


Figure S2: **Scalp distribution of the STAI-by-valence coefficient.** The map is descriptive. No channel survived FDR correction.

Theta gain was associated with NoGo-minus-Go performance ($\rho = 0.379$, permutation $p = 0.0098$, Holm-adjusted $p = 0.0294$, 95% bootstrap CI $[0.103, 0.613]$). Separate theta-gain associations with NoGo and Go performance were not supported after correction (Figure S3).

Theta gain and P3-minus-N2 gain were positively associated across participants. This participant-level covariation characterizes convergence between the measures but does not establish that they index the same neural process (Figure S4).

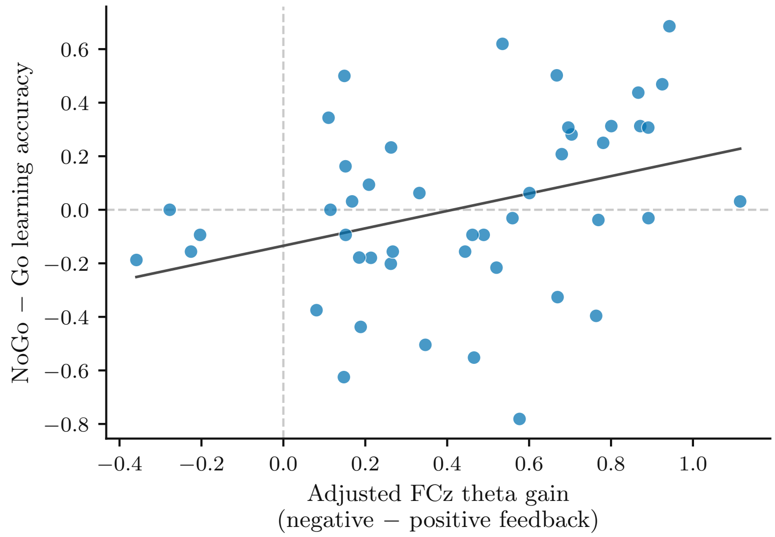


Figure S3: **Theta gain and NoGo-minus-Go learning.** Each point represents one elevated-BDI participant. The line is descriptive.

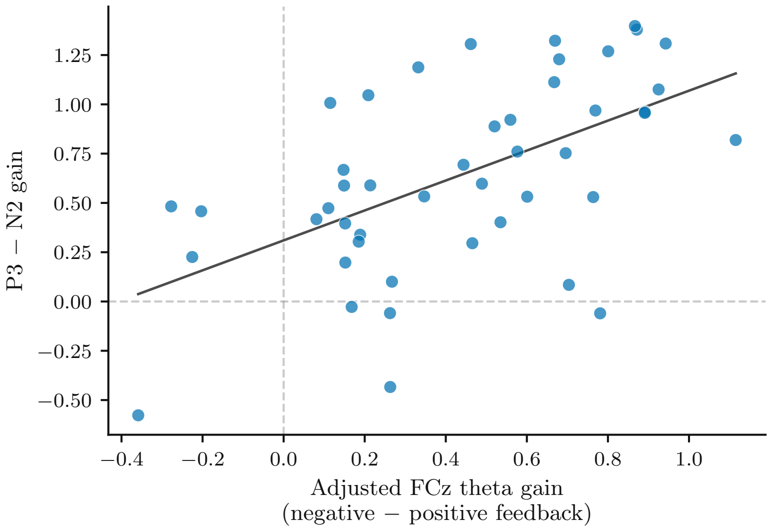


Figure S4: **Participant-level theta and P3-minus-N2 gain.** The line is descriptive.

#### 2.3 Components of signed-PE identification

The empirical within-participant distribution of signed PE by feedback valence is shown in Figure S13. In the rectified grid, target recovery was the across-participant correlation between generating and estimated neural slopes, averaged over replications. Target detection was the proportion of replications with a significant STAI slope. Coefficient estimability was assessed from each design matrix, and competing-predictor detection measured cross-attribution to the alternative adjusted regressor. These components were evaluated separately before applying the four-part criterion. Across the full simulation grid, signed-PE target recovery passed its criterion in 69.8% of cells, target detection in 74.6%, target-coefficient estimability in 100%, and the competing-predictor criterion in 43.7%. All four criteria were met in 40.1% of signed-PE cells. The corresponding four-part success rate for feedback valence was 96.8%. Figures S5–S7 show the recovery, target-detection, and competing-predictor surfaces under moderate observation noise. At high predictor overlap, attribution between correlated predictors failed more often than coefficient estimability.

#### 2.4 Independent electrophysiological analyses

In Study 1, the independent feedback response was strongest in the 200–400-ms interval relative to both flanking windows. Placebo showed the same temporal ordering, although the 200–400- versus 400–600-ms contrast was less precise (Figure S8).

P3-minus-N2 showed the same negative-minus-positive direction as theta in both independent cohorts (Figure S9). The shared direction is consistent with convergence between the conventional time-domain measure and the time–frequency response. The analysis was not designed as an equivalence test.

Detailed independent-dataset reliability estimates are shown in Figure S10. Study 1 estimates were high and comparatively precise. Placebo first–second estimates were uncertain because the eligible cohort was small.

#### 2.5 External computational and related-trait analyses

Partial $R^2$ values were small across all external joint models (Figure S11). Signed PE after valence accounted for the most incremental variance in one-rate Study 1, but even there the increment was small. Valence after signed PE accounted for more incremental variance in the dual-rate Study 1 model and in both placebo models. Descriptive trial-level Bayesian information criterion (BIC) values are reported in Supplementary Table S1.

Among 68 Study 1 participants with complete Behavioral Inhibition System (BIS) and BDI data, BIS was positively associated with negative-minus-positive FCz theta gain after adjustment for BDI (partial Spearman $\rho = 0.283$, permutation $p = 0.0218$, 95% bootstrap CI $[0.044, 0.498]$; Figure S12). BIS is reported as a related behavioral-inhibition measure and is not used as an STAI substitute.

Table S1: Descriptive trial-level BIC comparison of external encoding models.

| Cohort | PE model | ΔBIC | Descriptive BIC winner |
|---|---|---|---|
| Study 1 untreated | One-rate | -8.90 | Joint |
| Study 1 untreated | Dual-rate | 9.28 | Valence only |
| Placebo | One-rate | 6.68 | Valence only |
| Placebo | Dual-rate | 5.27 | Valence only |

*Note.* $\Delta\text{BIC} = \text{BIC}_{\text{joint}} - \text{BIC}_{\text{valence-only}}$. Negative values favor the joint model. These trial-level information criteria are descriptive because the likelihood does not represent within-session dependence. The main manuscript uses leave-one-session-out predictive comparison.

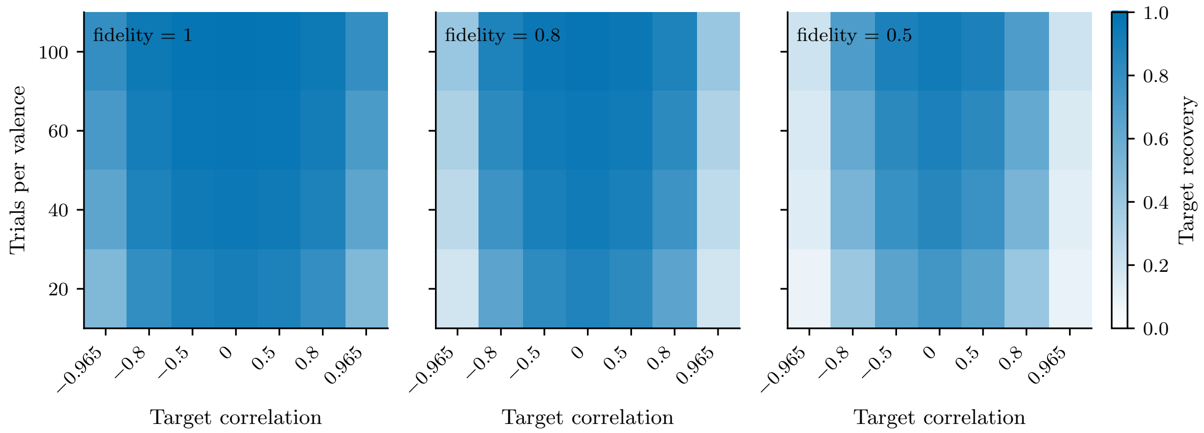


Figure S5: **Signed-PE target recovery under moderate noise.** Recovery of participant-specific generating neural slopes is shown across predictor overlap, estimated-PE fidelity, and trial count.

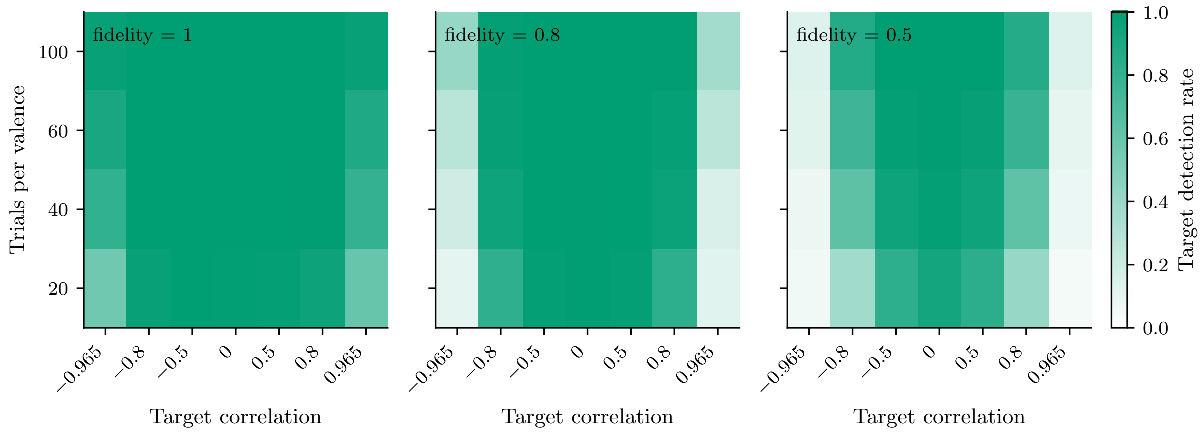


Figure S6: **Signed-PE target detection under moderate noise.** The proportion of replications detecting the target STAI association at $p < 0.05$ is shown across predictor overlap, estimated-PE fidelity, and trial count.

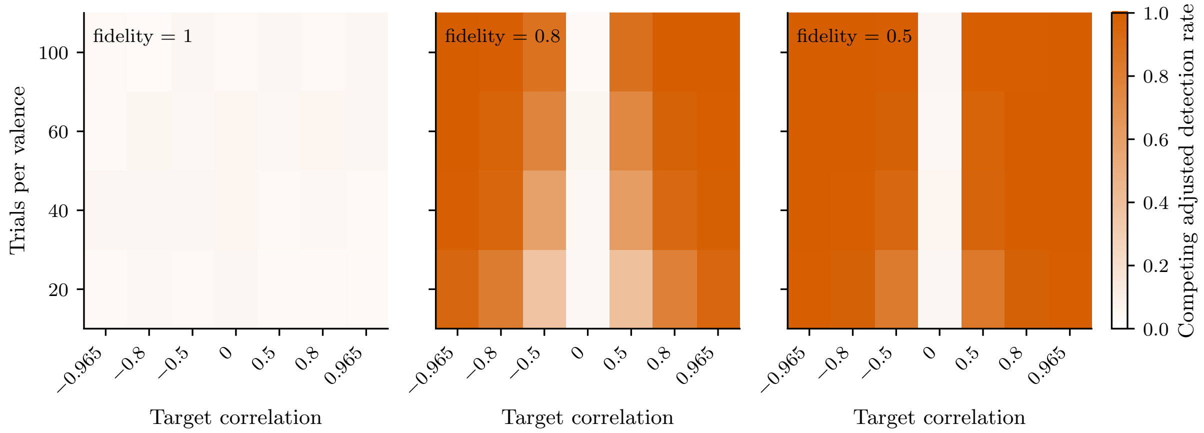


Figure S7: **Competing-predictor detection under moderate noise.** Values show detection of the competing adjusted predictor (cross-attribution) in signed-PE-generating simulations.

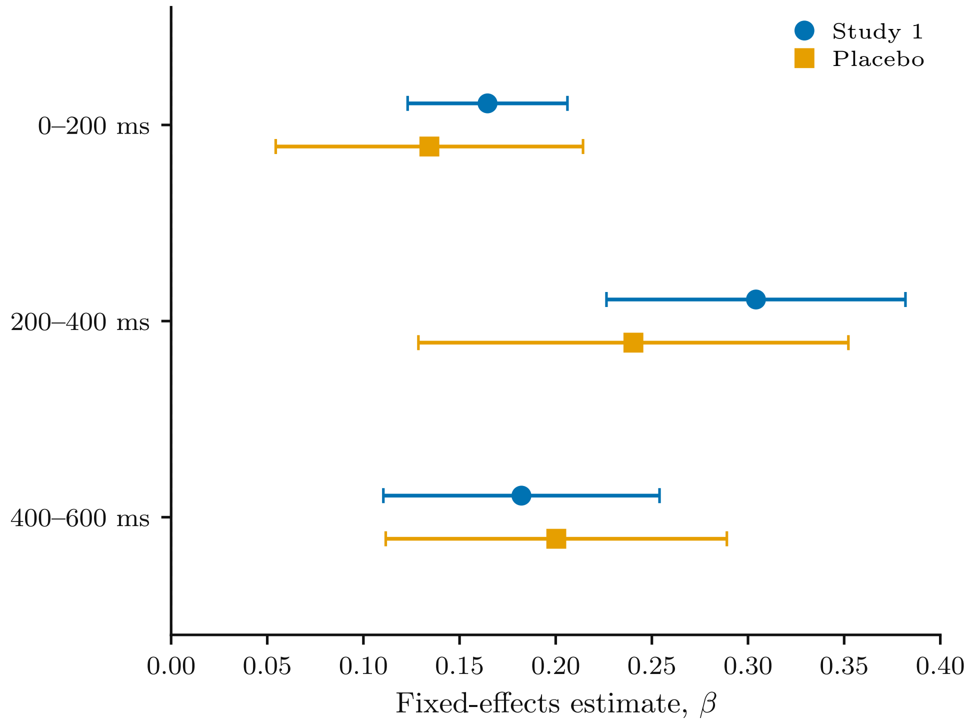


Figure S8: **Temporal profile of the independent feedback response.** Estimates are shown for Study 1 and placebo.

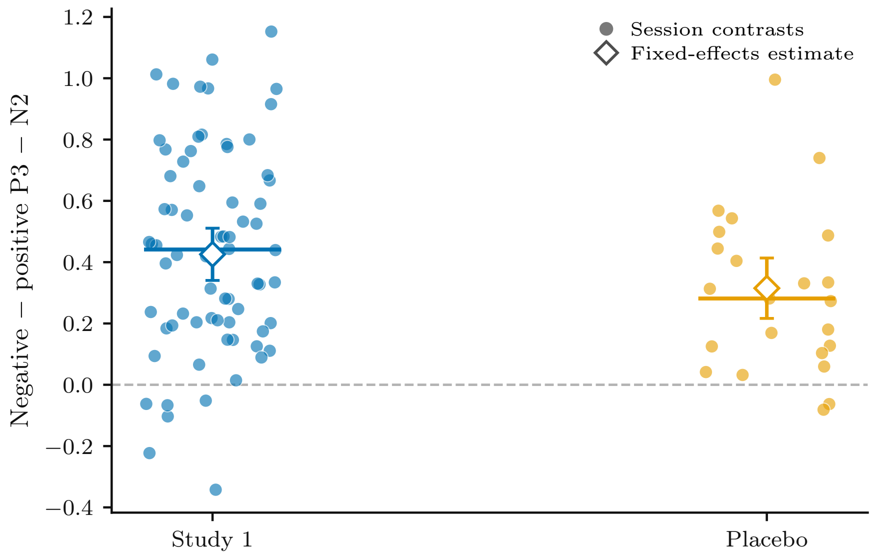


Figure S9: **Independent P3-minus-N2 feedback response.** Study 1 and placebo estimates are shown with confidence intervals.

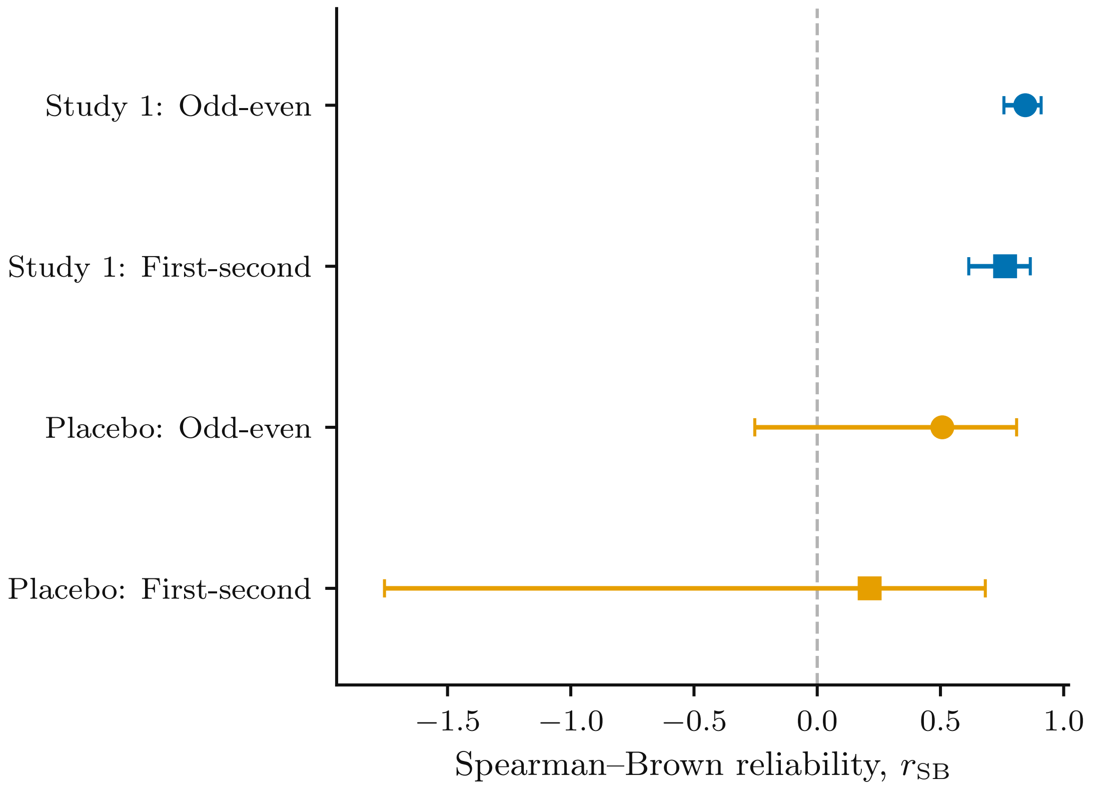


Figure S10: **Reliability of the independent feedback response.** Split-half estimates are shown for Study 1 and placebo.

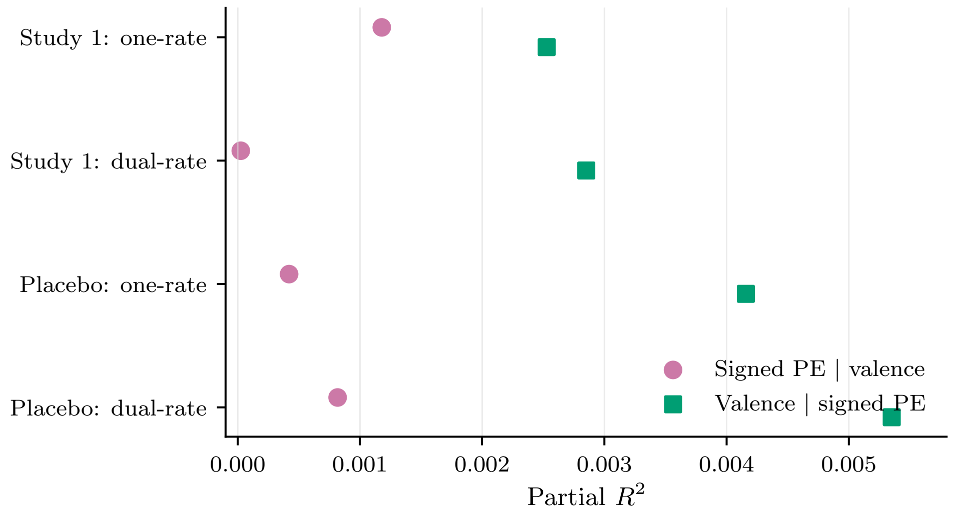


Figure S11: **Incremental variance in external joint models.** Partial $R^2$ is shown for signed PE after valence and valence after signed PE.

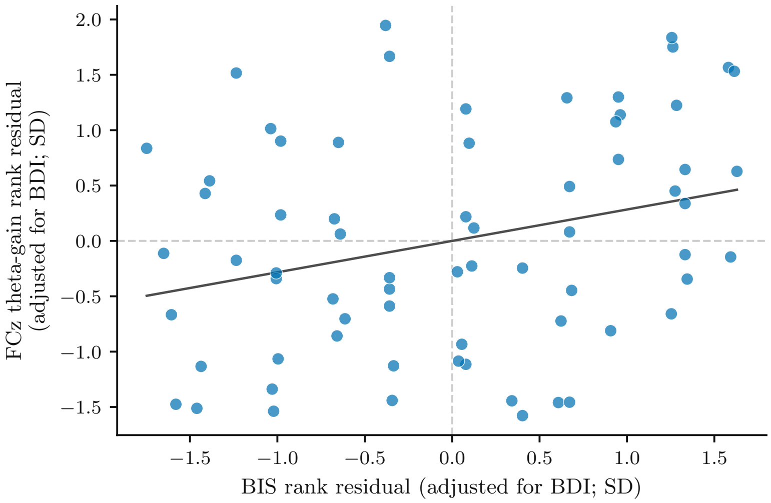


Figure S12: **BIS and theta gain after BDI adjustment.** Axes are standardized rank residuals. The line is descriptive.

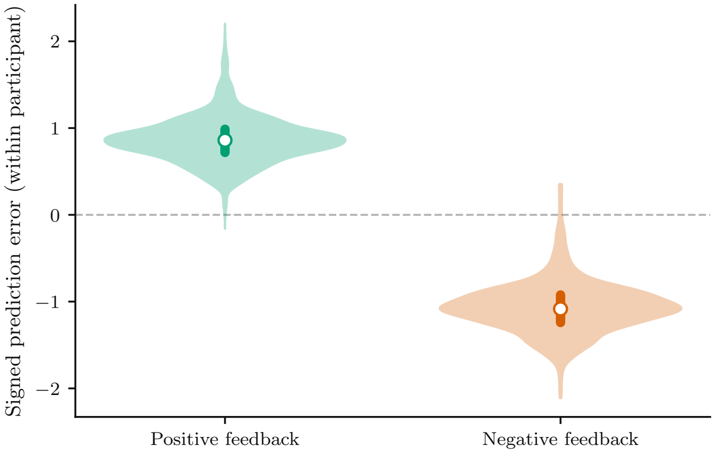


Figure S13: **Empirical feedback-valence and signed-PE overlap in the primary task.** Within-participant signed PE is shown separately for positive and negative feedback.
